# The nucleoid occlusion protein SlmA is a direct transcriptional activator of chitobiose utilization in *Vibrio cholerae*

**DOI:** 10.1101/141812

**Authors:** Catherine A. Klancher, Chelsea A. Hayes, Ankur B. Dalia

## Abstract

Chitin utilization by the cholera pathogen *Vibrio cholerae* is required for its persistence and evolution via horizontal gene transfer in the marine environment. Genes involved in the uptake and catabolism of the chitin disaccharide chitobiose are encoded by the *chb* operon. The orphan sensor kinase ChiS is critical for regulation of this locus, however, the mechanisms downstream of ChiS activation that result in expression of the *chb* operon are poorly understood. Using an unbiased transposon mutant screen, we uncover that the nucleoid occlusion protein SlmA is a regulator of the *chb* operon. SlmA has not previously been implicated in gene regulation. Also, SlmA is a member of the TetR family of proteins, which are generally transcriptional repressors. *In vitro*, we find that SlmA binds directly to the *chb* operon promoter, and *in vivo*, we show that this interaction is, surprisingly, required for transcriptional activation of this locus and for chitobiose utilization. Using point mutations that disrupt distinct functions of SlmA, we find that DNA-binding, but not nucleoid occlusion, is critical for transcriptional activation. This study identifies a novel role for SlmA as a transcriptional regulator in *V. cholerae* in addition to its established role as a cell division licensing factor.

**AUTHOR SUMMARY:** The cholera pathogen *Vibrio cholerae* is a natural resident of the aquatic environment and causes disease when ingested in the form of contaminated food or drinking water. In the aquatic environment, the shells of marine zooplankton, which are primarily composed of chitin, serve as an important food source for this pathogen. The genes required for the utilization of chitin are tightly regulated in *V. cholerae*, however, the exact mechanism underlying this regulation is currently unclear. Here, we uncover that a protein involved in regulating cell division is also important for regulating the genes involved in chitin utilization. This is a newly identified property for this cell division protein and the significance of a common regulator for these two disparate activities remains to be understood.

## INTRODUCTION

*V. cholerae*, the bacterium responsible for the diarrheal illness cholera, is naturally found in the marine environment. In this niche, this pathogen form biofilms on the shells of microscopic crustaceans. The shells of these organisms are primarily composed of chitin, an insoluble polymer of β−1,4 linked *N*-acetylglucosamine (GlcNAc). Chitin is the second most abundant biopolymer on the planet, and *Vibrio* species play an important role in recycling chitin in the aquatic environment [1]. The pathway for degradation and utilization of this carbon and nitrogen source is conserved among the *Vibrionaceae* [2].

In addition to promoting the survival of *V. cholerae* in the aquatic environment, biofilm formation on chitin is also important for promoting transmission of this pathogen to its human host. Indeed, cholera outbreaks are seasonal in endemic areas and closely associated with blooms in chitinous zooplankton [1]. Furthermore, filtration of water through sari cloth in these areas (which has an effective pore size that will eliminate chitin biofilms but cannot filter planktonic bacteria) reduces the incidence of waterborne transmission [3, 4]. Chitin oligosaccharides also induce natural competence, a mechanism of horizontal gene transfer, in *V. cholerae* [5, 6]. Thus, *Vibrio-chitin* interactions are important for the persistence, transmission, and evolution of this important human pathogen in its environmental reservoir.

The expression of genes required for chitin degradation, uptake, and utilization are regulated by the orphan sensor kinase ChiS [7, 8]. ChiS activity is normally repressed by chitin binding protein (CBP), the periplasmic substrate binding protein for the chitobiose ABC transporter [7]. This repression is relieved in the presence of chitin oligosaccharides, which bind to CBP. ChiS can also be activated, however, by deletion of CBP [7]. ChiS is an orphan sensor kinase and is not predicted to directly bind DNA. Without a cognate response regulator, the mechanism of gene regulation downstream of ChiS activation is poorly understood. One locus regulated by ChiS is the *chb* operon, which encodes the genes required for uptake and utilization of the chitin disaccharide chitobiose [8]. Here, we perform an unbiased screen to identify factors downstream of ChiS required for activation of the *chb* operon and characterize one newly identified transcriptional activator of this locus.

## RESULTS

### An unbiased screen identifies SlmA as a putative activator of chitin utilization genes in V. cholerae

To study ChiS regulation of the *chb* operon, we first generated a P_*chb*_ transcriptional reporter. As expected, this reporter was activated in a CBP mutant, and this activation was dependent on ChiS (**Fig. 1A**). We exploited this fact and used our transcriptional reporter to perform two independent transposon mutant screens to identify potential activators and repressors of this locus. To identify repressors, we used a P_*chb*_-*lacZ* reporter (which forms white colonies), and screened for mutant blue colonies. Conversely, we screened for activators by using a P_*chb*_-*lacZ* Δ*cbp* strain (which forms blue colonies) and isolated mutant white colonies. A positive control for the repressor screen was *cbp*, while a positive control for the activator screen was *chiS*. As expected, both of these genes were identified in their respective screens, which helped to validate this approach. No additional hits were identified in the repressor screen, but one hit that we identified in the activator screen was VC0214, the gene encoding the nucleoid occlusion protein SlmA (**Fig. 1A**). Consistent with a role in transcriptional activation of this locus, P_*chb*_ is no longer induced in a Δ*slmA* Δ*cbp* mutant strain (Fig. 1A and B). To determine if SlmA is also required for activation of this locus under more physiologically relevant conditions, we assessed *cbp* transcript levels in WT and Δ*slmA* cells when induced with (GlcNAc)_6_, the chitin hexasaccharide. While this locus is strongly induced in the WT, there is little to no activation in the *slmA* mutant, which is consistent with what we observed when using a *cbp* mutant to artificially activate this locus (**Fig. 1D**). Using 5’ RACE, we mapped the transcription start sites (TSS) for the *chb* transcript and found that there are two distinct TSSs for this locus. One TSS representing a longer transcript that is basally expressed in the absence of chitin induction (uninduced transcript), and a second TSS for a shorter transcript that is expressed when this locus is induced by chitin oligosaccharides or artificially induced by deletion of CBP (induced transcript) (**Fig. S1**). Using primers that are specific to the longer uninduced transcript, we find that levels of this transcript are induced only 2 to 4-fold when cells are grown in the presence or absence of (GlcNAc)_6_ in both WT and Δ*slmA* cells, indicating that SlmA is specifically playing a role in the activation of this locus at the downstream TSS (**Fig. S1**).

**Figure 1.**
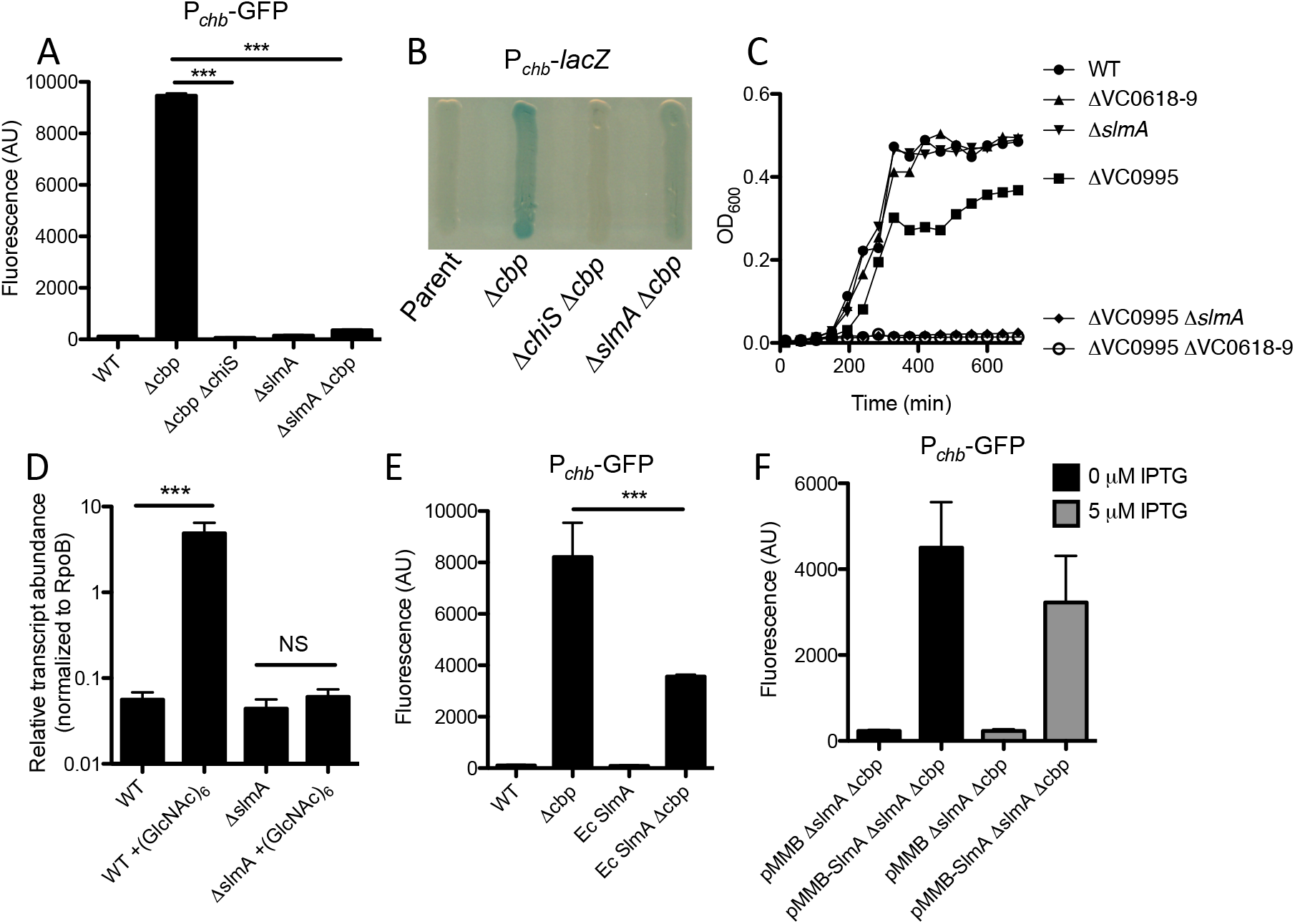
-SlmA is required for transcriptional activation of *P_chb_*. (**A**) GFP fluorescence was measured in the indicated strains, all of which contain a P_*chb*_-*gfp* transcriptional reporter. Statistics shown indicate that samples are significantly different from the “Δ*cbp*” sample. (**B**) Transcription of P_*chb*_ was schematized on X-gal-containing media in the indicated strains, all of which contained a P_*chb*_-*lacZ* transcriptional reporter. (**C**) Growth curves of the indicated strains in M9 minimal medium containing 0.5% chitobiose as a sole carbon source. (**D**) qRT-PCR of the *chb* short transcript was measured in WT and Δ*slmA* cells grown in the presence or absence of (GlcNAc)6. (**E**) GFP fluorescence was measured in the indicated strains, all of which contain a P_*chb*_-*gfp* transcriptional reporter. (**F**) GFP fluorescence was measured in a P_*chb*_-*gfp* Δ*slmA* Δ*cbp* strain harboring either an empty vector (pMMB) or a SlmA expression vector (pMMB-SlmA) grown in the presence or absence of 5 μM IPTG. Data in **A**, **D**, **E**, and **F** are shown as the mean ± SD and are from at least three independent biological replicates. Data from **B** and **C** are representative of at least 2 independent experiments. *** = p<0.001, NS = not significant.

The *chb* operon encodes an ABC transporter that mediates uptake of chitobiose (permease encoded by VC0618-0619). *V. cholerae*, however, can also degrade chitobiose in the periplasm into the chitin monomer GlcNAc [2, 9, 10], which is taken up by a PTS transporter, VC0995, that is outside of the P_*chb*_ operon and whose regulation is independent of ChiS (**Fig. S2**) [8]. Thus, deletion of either transporter independently does not eliminate growth on chitobiose (**Fig. 1C**). However, upon deletion of both the chitobiose and GlcNAc transporters, cells are unable to grow on chitobiose as the sole carbon source (**Fig. 1C**). To determine if SlmA-dependent gene regulation of P_*chb*_ was physiologically relevant, we assessed whether SlmA was required for growth on chitobiose ((GlcNAc)_**2**_) as a sole carbon source. When we performed this analysis, we found that SlmA was required for growth on chitobiose in a ΔVC0995 mutant background, which is consistent with SlmA playing a physiologically important role in regulating the chitobiose transporter in the *chb* operon. As expected, all strains grew equally well when glucose or tryptone was provided as the sole carbon source whereas any strains with a ΔVC0995 mutation were unable to grow on GlcNAc (**Fig. S2**). A *slmA* mutant, however, does not phenocopy a *chiS* mutant. In fact, a Δ*chiS* strain is unable to grow on (GlcNAc)_2_ as the sole carbon source, which is not surprising since this regulator is required for the expression of a number of chitin catabolic genes in addition to the *chb* operon [8](**Fig. S3**).

SlmA function has primarily been characterized in *E. coli* [11–15]. This protein, however, is well conserved between *E. coli* and *V. cholerae* (67% identity, 83% similarity; **Fig. S4**). To determine if gene regulation by SlmA is a conserved property, we replaced the native copy of *slmA* in *V. cholerae* with the gene for *slmA* from *E. coli*. We found that *Ec* SlmA was able to activate transcription of P_*chb*_, albeit, less robustly than *Vc* SlmA, suggesting that this activity is not unique to the *Vc* homolog, but is a conserved property of this protein (**Fig. 1E**).

To further confirm that deletion of *slmA* was responsible for the phenotypes observed on the P_*chb*_ operon, we complemented *slmA* on a plasmid. As expected, we found that ectopic expression of SlmA was sufficient to recover activation of P_*chb*_ in a Δ*slmA* Δ*cbp* mutant (**Fig. 1F**). As seen previously [16], expression from the pMMB vector used here is leaky and as a result, basal expression of SlmA from this plasmid without inducer is sufficient to activate P_*chb*_ (**Fig. 1F**).

### SlmA is a direct transcriptional activator of Pchb

SlmA is a TetR family protein and binds to a specific and well-defined sequence [12]. Through bioinformatic analysis, we identified one SlmA binding site (SBS) in P_*chb*_ based on the SBS consensus sequence of *E. coli* SlmA (**Fig. 2A**) [12]. This was further confirmed by a recent *in vitro* whole genome binding analysis that identified putative SBSs in *V. cholerae* [17].

**Figure 2.**
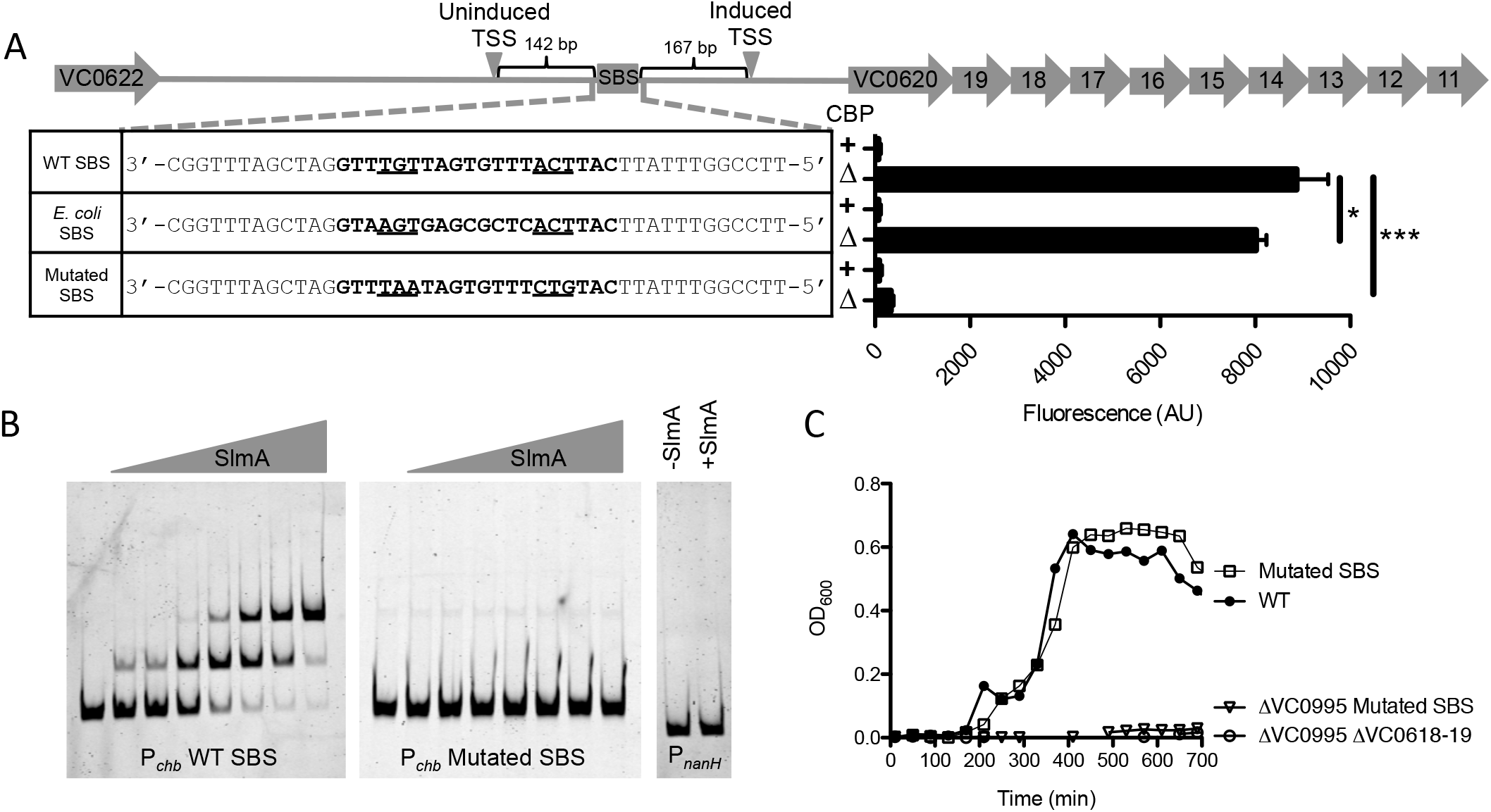
-SlmA is a direct transcriptional activator of P_chb_. (**A**) Promoter map of P_*chb*_ showing the SBS and the TSSs mapped in the presence (induced) and absence (uninduced) of chitin. “WT SBS” represents the native P_*chb*_ sequence, “*E. coli* SBS” indicates that the native SBS was swapped for the consensus SBS from *E. coli*, and “mutated SBS” indicates that the 6 highly conserved residues in the SBS were mutated. SBS sequences for the indicated mutants are shown in bold text and the 6 most highly conserved residues of the SBS are underlined. GFP fluorescence was measured in strains harboring the indicated mutations in the P_*chb*_-*gfp* transcriptional reporter. The *cbp* status of each strain is indicated by a “+” (*cbp* intact) or a “Δ” *(Δcbp* strain background). Data are shown as the mean ± SD and are from at least three independent biological replicates. (**B**) EMSAs performed with purified SlmA and the indicated promoter probes. The two P_*chb*_ probes were incubated with (from left to right) 0 nM, 7.5 nM, 15 nM, 30 nM, 60 nM, 120 nM, 240 nM, or 480 nM SlmA. The P_*nanH*_ probe was incubated with 0 nM or 480 nM SlmA. (**C**) Growth curves of the indicated strains in M9 minimal medium containing 0.5% chitobiose as the sole carbon source. Data from **B** and **C** are representative of at least 2 independent experiments. *** = *p*<0.001, * = p<0.05, NS = not significant.

To determine if SlmA could bind to this site, we performed electrophoretic mobility shift assays (EMSAs) using a probe from P_*chb*_ that contains the putative SBS. Indeed, we found that SlmA could directly bind to P_*chb*_ (**Fig. 2B**). SlmA was previously shown to bind to an SBS as a dimer-of-dimers [14], which is consistent with the two shifts we observed with the P_*chb*_ promoter probe (**Fig. 2B**). As a negative control, we tested binding of SlmA to *PnanH*, an unrelated promoter that lacks a predicted SBS. As expected, we found that SlmA could not bind to this probe (**Fig. 2B**). To determine if the putative SBS in P_*chb*_ is responsible for the shift observed in our EMSAs, we mutated 6 highly conserved residues in the SBS. We found that SlmA no longer shifted the P_*chb*_ probe when these residues were mutated, suggesting that this sequence represents a *bona fide* SBS in the *chb* promoter (**Fig. 2B**).

Since we found that SlmA could directly bind to an SBS in the *chb* promoter, we next wanted to determine if this site is important for the transcriptional activation of this locus. To that end, we assessed activation of P_*chb*_ in a mutant strain where we mutated the SBS. When the SBS was mutated, we no longer observed activation of P_*chb*_ (**Fig. 2A**). Promoter truncations of P_*chb*_ also confirmed that the SBS in this promoter was required for activation of this locus (**Fig. S5**). Furthermore, we replaced the native SBS in the P_*chb*_ promoter with the consensus SBS for *E. coli* SlmA. This alternative SBS was able to support activation of the locus (**Fig. 2A**). Cumulatively, these results suggest that SlmA recruitment to the P_*chb*_ promoter via an SBS is required for transcriptional activation. Our previous results indicated that *Ec* SlmA supported activation of P_*chb*_ poorly compared to *Vc* SlmA (**Fig. 1E**). To determine if this was due to a reduced affinity of *Ec* SlmA for the SBS in P_*chb*_, we tested activation of P_*chb*_ containing the consensus *Ec* SBS in a strain expressing *Ec* SlmA at the native locus. This strain, however, had similar levels of P_*chb*_ induction compared to a strain with *Ec* SlmA and the native SBS at P_*chb*_ (**Fig. S6**). Thus, reduced activation of Ec SlmA is likely due to a reduced ability to promote transcriptional activation and not due to reduced affinity for the SBS.

Above, we showed that SlmA was required for regulation of the chitobiose ABC transporter encoded by the *chb* operon for optimal growth on chitobiose (**Fig. 1C**). Deletion of SlmA, however, may have pleiotropic effects. To determine if SlmA binding at the SBS in P_*chb*_ was responsible for this effect, we mutated the SBS in the native *chb* promoter and tested growth on chitobiose. We found that mutating the SBS in P*chb* prevents growth on chitobiose in the ΔVC0995 background, which is consistent with this SBS being critical for regulating the chitobiose transporter in the *chb* operon (**Fig. 2C**). These results indicate that SlmA-dependent regulation of P_*chb*_ is both direct and physiologically relevant.

### DNA binding, but not nucleoid occlusion activity, is required for SlmA-dependent transcriptional activation of Pchb

The ability of SlmA to mediate nucleoid occlusion is dependent upon its ability to dimerize, bind DNA, and interact with FtsZ. The residues involved in these interactions have been previously characterized [11–14]. So, we next wanted to dissect which functional interactions of SlmA are required for transcriptional activation of P_*chb*_. To that end, we mutated residues in SlmA involved in DNA binding (T31, E43), FtsZ interaction (F63, R71), and dimerization (R173) (**Fig. 3A**). We then tested these mutants for their ability to activate expression of P_*chb*_. We also assessed whether these SlmA variants could bind to DNA using a previously described synthetic SBS-*gfp* reporter where SlmA binding at a high affinity SBS represses expression of GFP [11]. We used the consensus SBS from *E. coli* instead of the native SBS found in P_*chb*_ because SlmA binds to this SBS with a higher affinity, allowing for a more sensitive evaluation of DNA binding (**Fig. S7**). We also tested the nucleoid occlusion activity of these SlmA variants by overexpressing each and assessing the morphology of cells. Overexpression of WT SlmA results in a dramatic filamentous phenotype due to excess nucleoid occlusion activity, whereas alleles deficient in nucleoid occlusion do not cause this phenotype [15](**Fig. S5**). Also, we triple-FLAG tagged all SlmA alleles tested at the native locus to assess their expression levels by western blot analysis, which uncovered that all were expressed at least at WT levels (**Fig. S8**). Importantly, SlmA-triple FLAG constructs were still functional for P_*chb*_ activation (**Fig. S8**).

**Figure 3.**
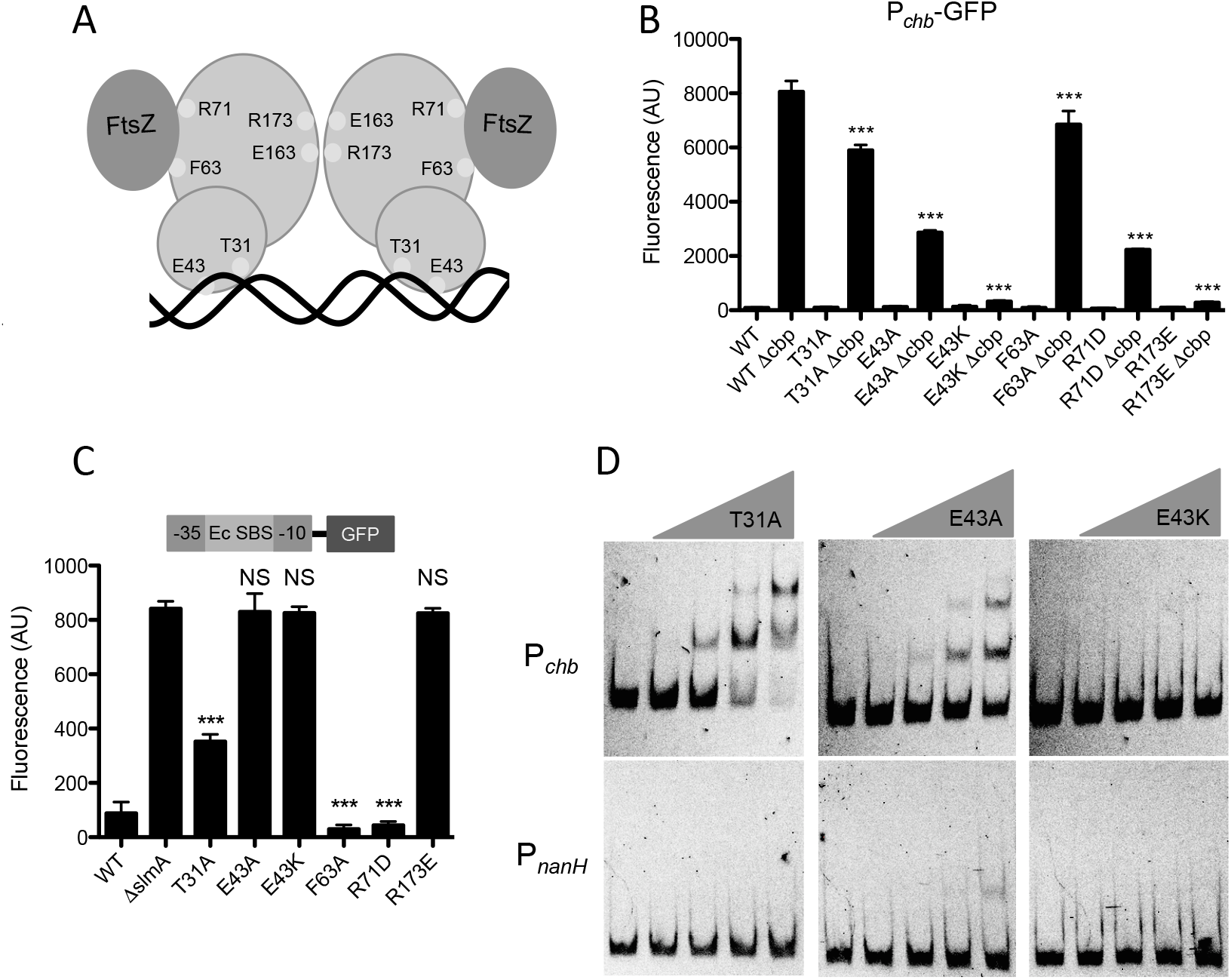
-DNA binding, but not nucleoid occlusion activity, is required for SlmA-dependent transcriptional activation of  *P_chb_*. (**A**) Schematic of SlmA depicting residues mutated in this study. T31 and E43 are predicted to interact with DNA, F63 and R71 mediate FtsZ interactions, and R173 is required for dimerization. (**B**) GFP fluorescence was measured in strains containing a P_*chb*_-*gfp* reporter and the indicated site-directed mutations in the native copy of *slmA*. Statistical comparisons are made to the “WT Δ*cbp*” sample. (**C**) GFP fluorescence was assessed in strains harboring an SBS-*gfp* reporter that is a readout for DNA binding and contain the indicated site-directed mutations in the native copy of *slmA*. Statistical comparisons are made to the “Δ*slmA*” sample. (D) EMSAs performed with the WT P_*chb*_ probe or the negative control P_*nanH*_ probe incubated with (from left to right): 0 nM, 240 nM, 480 nM, 960 nM, or 1920 nM of the indicated SlmA mutant protein. Data in **B** and **C** are shown as the mean ± SD and are from at least three independent biological replicates. Data from **D** is representative of at least two independent experiments. *** = *p*<0.001, NS = not significant.

First, we found that the FtsZ binding mutants (F63A and R71D) were still able to facilitate transcriptional activation of P_*chb*_, although one mutant (R71D) displayed a reduced degree of activation (**Fig. 3B**). As expected, these SlmA variants were still able to bind DNA (**Fig. 3C**) and had lost nucleoid occlusion activity (**Fig. S9**). These results suggest that the nucleoid occlusion activity of SlmA is not required for its ability to act as a transcriptional activator at the *chb* operon.

Next, we tested DNA binding mutants T31A and E43A that were previously identified in *E. coli* [11], and, surprisingly, we found that these variants were still able to activate transcription of P_*chb*_. The T31A mutant was still able to bind DNA as indicated by the synthetic SBS-GFP reporter (**Fig. 3C**). The E43A mutant, however, was not able to bind DNA in this assay (**Fig. 3C**). Our previous results with the mutated SBS would suggest that SlmA binding to P_*chb*_ is required for activation, therefore, we hypothesized that the E43A mutant may still have the ability to bind an SBS, however, this may be below the limit of detection of our DNA binding reporter. To test this, we purified SlmA E43A and assessed its ability to bind P_*chb*_ in an EMSA. Indeed, we found that SlmA E43A was still able to bind to this promoter, although with a greatly reduced affinity compared to WT SlmA or SlmA T31A (**Fig. 3D** and **2B**). Thus, these results indicate that while these mutations may eliminate DNA binding of Ec SlmA [11], they may have a limited or reduced impact on DNA-binding of Vc SlmA. Overexpression of these DNA binding mutants indicated that they had lost the ability to mediate nucleoid occlusion (**Fig. S9**).

Dimerization of SlmA is required for DNA binding as well as nucleoid occlusion [11]. Dimerization requires a charge-based interaction of R173 from one monomer with E163 of the other monomer. Swapping the charge of either residue can prevent dimerization [11], thus we used an R173E mutant to test the role of SlmA dimerization. The dimerization mutant R173E was unable to activate P_*chb*_. As expected, this variant had also lost the ability to bind DNA (**Fig. 3C**) and mediate nucleoid occlusion (**Fig. S9**).

Next, we sought to identify residues that are required for SlmA-mediated activation of P_*chb*_. To that end, we performed error-prone PCR of SlmA and screened for colonies that had lost the ability to activate P_*chb*_-*lacZ*. We then counter-screened colonies for the ability to bind DNA using our synthetic SBS-GFP reporter. We screened >15,000 colonies and identified ~500 colonies that were deficient for activation of P_*chb*_-*lacZ*. In our counter-screen, however, we found that none of these SlmA mutants maintained the ability to bind DNA, suggesting that DNA binding and transcriptional activation of P_*chb*_ are tightly linked. One variant identified in this screen was SlmA E43K, which is the same residue previously identified as playing a role in DNA binding [14], however, with a lysine substitution in place of alanine. We hypothesized that the E43K mutant could no longer bind to DNA, in contrast to the E43A variant studied above. To test this, we purified SlmA E43K and assessed its ability to bind P_*chb*_ *in vitro*. Indeed, by EMSA we found that E43K could not bind P_*chb*_ (**Fig. 3D**). Importantly, this mutation does not simply result in an unstable protein because the E43K mutant still makes WT levels of SlmA protein (**Fig. S8**). This mutant also lost the ability to mediate nucleoid occlusion when overexpressed (**Fig. S9**). Cumulatively, these results indicate that high affinity binding of SlmA to DNA may be required for proper nucleoid occlusion, while weak binding is sufficient to mediate transcriptional activation of P_*chb*_.

### SlmA is not sufficient to promote activation of P_*chb*_

Thus far, our data suggest that activation of P_*chb*_ requires activation of ChiS, either via deletion of *cbp* or induction with chitin oligosaccharides. To determine if SlmA is the sole activator that acts downstream of ChiS we decided to overexpress SlmA to see if it was sufficient to activate P_*chb*_ in the absence of either inducer. Overexpression of WT SlmA results in a dramatic filamentous phenotype (**Fig. S9**). Interestingly, overexpression of WT SlmA from a chromosomally integrated Ptac-SlmA construct still induced P_*chb*_ expression in a Δ*cbp*-dependent manner, even at levels that induced morphological defects (10 μM), although activation was lost when cultures were induced with 100 μM IPTG (**Fig. S10**). Above, we show that the nucleoid occlusion mutants T31A, F63A, and R71D are still able to activate P_*chb*_ (**Fig. 3B**), while their overexpression does not result in filamentation (**Fig. S9**). Indeed, overexpression of SlmA F63A from a chromosomally integrated Ptac-SlmA F63A construct still supported activation in a Δ*cbp*-dependent manner, however, induction with 100 μM IPTG still prevented activation similar to what was observed with overexpression of WT SlmA (**Fig S10**). Also, this was independent of any obvious morphological defect, suggesting that overexpression of SlmA at this level prevents P_*chb*_ activation independent of its role in nucleoid occlusion (**Fig. S10**).

Next, To determine if SlmA was sufficient to mediate P_*chb*_ activation, we overexpressed SlmA T31A, F63A, and R71D mutants with 10 μM IPTG and assessed whether we could activate P_*chb*_ under non-inducing conditions (i.e. in a *cbp*^+^ strain grown in the absence of chitin oligosaccharides). Overexpression of these constructs, however, did not result in activation under non-inducing conditions (**Fig. S11**). Cumulatively, these results indicate that the repression of ChiS by CBP plays a dominant role over the activity of SlmA as a transcriptional activator of P_*chb*_. This is consistent with a model whereby another, as yet unidentified, protein acts downstream of ChiS and is required for coactivation of this locus along with SlmA.

Since overexpression of SlmA F63A with 100 μM IPTG inhibited P_*chb*_ activation, we decided to test whether this phenotype was the result of excess SlmA protein, which titrated away a putative co-activator from the P_*chb*_ operon. To that end, we overexpressed ChiS, which might enhance the expression / activity of this putative downstream coactivator. Interestingly, ChiS overexpression at 10–1000 μM also prevented P_*chb*_ activation, albeit, not to the degree of SlmA F63A overexpression (**Fig. S10**). ChiS overexpression, however, did not rescue the loss of P_*chb*_ activation when SlmA F63A was overexpressed (**Fig. S10**). The lack of increased P_*chb*_ expression over the native levels in the presence of higher levels of ChiS and/or SlmA indicate that neither protein is limiting and that native levels of these proteins are sufficient for maximal activation of P_*chb*_.

### Sequence specificity between the SBS and induced promoter for Pchb activation suggests a possible coactivator-binding site

The transcriptional regulator cAMP receptor protein (CRP) binds to a specific DNA sequence in the presence of cAMP, allowing for activation or repression of target genes.

One turn of B DNA is approximately 10 bp, and moving the CRP binding site in denominations of 10 bp proximally or distally from the promoter maintains transcriptional activation of genes regulated by CRP by maintaining the helical phase of the CRP binding site in relation to the promoter [18, 19]. Conversely, moving the CRP binding site in increments of less than 10 bp places the CRP binding site out of helical phase with the promoter and therefore prevents activation by disrupting proper interaction with RNA polymerase (RNAP).

We tested whether there was helical phase-dependence for SlmA-mediated activation of P_*chb*_ by a similar mechanism. We found that moving the SBS as little at 3 bp or as much as 12 bp resulted in a significant decrease in P_*chb*_ activation (**Fig. S12**). Constructs in which the SBS was moved to maintain the helical phase (i.e. in increments of 10 bp) were also unable to promote robust activation, indicating that SlmA may not activate transcription by direct contact with RNA polymerase in a mechanism similar to CRP (**Fig. S12**). This is in line with the distance of the SBS from the TSS, 167 bp, which makes it unlikely that SlmA directly interacts with RNAP to activate transcription. This supports the hypothesis that a coactivator may bridge SlmA and RNAP to mediate activation of this locus.

Next, we assessed whether there was sequence specificity for activation of P_*chb*_ in the region between the SBS and promoter element for the chitin-induced TSS. We hypothesized that if a coactivator was required for expression, there may be sequence specificity within this region, while if no coactivator was required, maintaining the spacing between the SBS and TSS may be sufficient to support activation. We tested this by replacing the region between the SBS and the −35 signal with the same number of nucleotides from an intergenic region of the *lacZ* gene (**Fig. 4**). The activation of this reporter was abolished after swapping the native sequence with *lacZ* sequence (**Fig. 4**), indicating that sequence specificity within this region may be required for activation. Furthermore, truncation studies indicated that elements necessary for activation include the SBS, TSS and intervening regions (**Fig. S5**). Also, a bioinformatically determined hairpin exhibits some inhibitory activity (**Fig. S5**). Together, these data are consistent with a model whereby the region between the SBS and TSS contains a coactivator-binding site required for P*chb* induction. There are, however, a number of alternative models that can account for the results obtained, which are addressed in the **Discussion** below.

**Figure 4.**
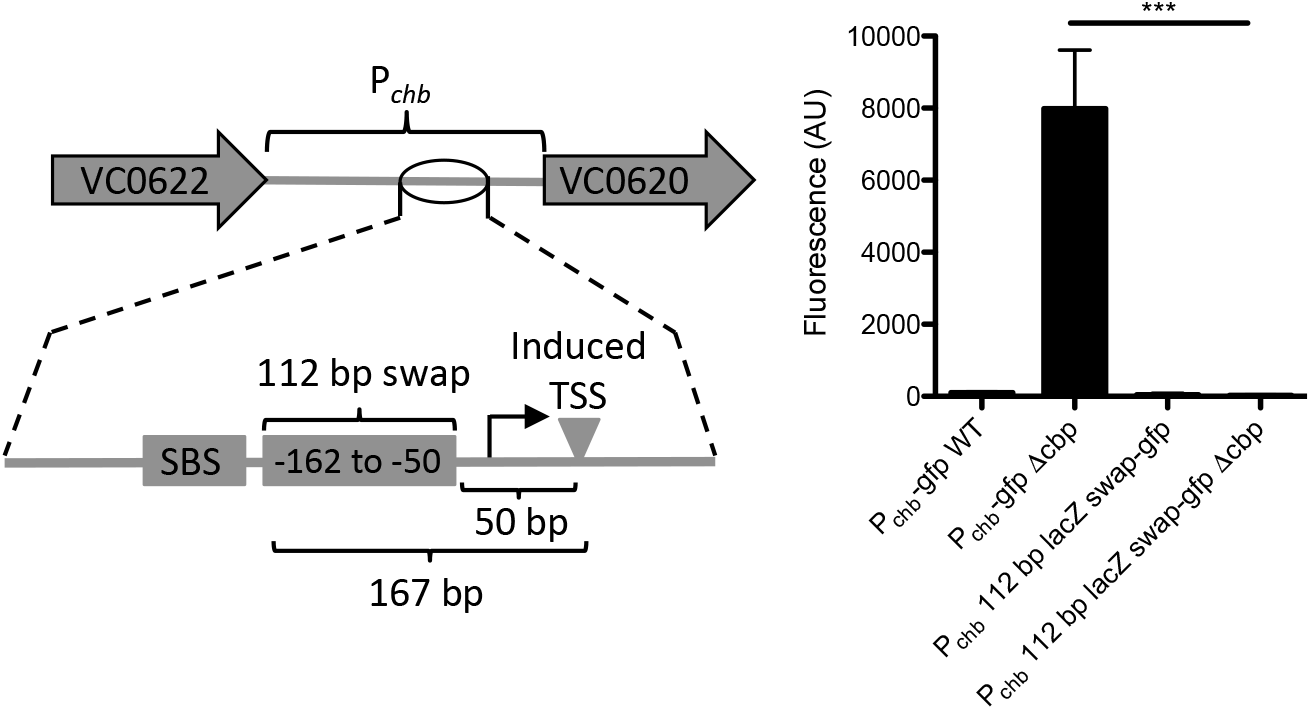
–Sequence specificity between the SBS and induced promoter for *P_chb_* activation. A 112 bp fragment of P_*chb*_-*gfp* (from −162 to −50 bp) was swapped out with a fragment of equal size from the *lacZ* gene and activation was determined by assessing fluorescence. All data are shown as the mean ± SD and are from at least three independent biological replicates.

Finally, we were interested in determining which sigma factor was required for activation of P_*chb*_. To test this, we inactivated every non-essential sigma factor (RpoN, RpoS, RpoE, RpoF, and RpoH) in *V. cholerae* and assessed transcriptional activation of P_*chb*_. We observed activation in all mutant strains, suggesting that the essential housekeeping sigma factor RpoD likely plays a dominant role in P_*chb*_ expression (**Fig. S13**). The degree of P_*chb*_ activation, however, was reduced in the *rpoE* mutant, suggesting that this sigma factor either plays a minor role in activation of this locus or that RpoE is required for proper expression or folding of the GFP reporter. Additionally, we observed an increase in expression in the *rpoS* mutant, suggesting that this sigma factor may play a role in the expression of some upstream repressor of this locus.

## DISCUSSION

Here, we show that the cell division licensing factor SlmA is required for activation of chitobiose utilization genes in *V. cholerae*. We show that this regulation is direct because SlmA binds to P_*chb*_, the promoter that regulates these genes. Furthermore, we demonstrate that this direct regulation is required for growth of *V. cholerae* on chitobiose.

While we show that SlmA is a direct activator of P_*chb*_, the exact mechanism of activation is still unclear. We have shown that the location of the SBS within P_*chb*_ is important for activation. Additionally, we show that the sequence between the SBS and the −35 sequence is required for activation, consistent with this region harboring a coactivator binding site. Because the SBS is located 167 base pairs away from the TSS, it is unlikely that SlmA directly interacts with RNA Polymerase to mediate activation. We also explored the possibility that SlmA binding may occlude a repressor binding site thereby allowing for activation of P_*chb*_ indirectly. We formally tested this hypothesis by performing a Tn mutant screen for repressors in a Δ*slmA* Δ*cbp* P_*chb*_-*lacZ* reporter strain, however, this screen did not identify any putative repressors (~70,000 mutants visually screened). As a result, we propose a mechanism in which SlmA requires a presently unknown ChiS-dependent coactivator for activation of P_*chb*_ (**Fig. 5**). A candidate for this unknown factor was not identified in the Tn-mutant screen for activators that identified SlmA. This screen, however, may not have been saturating or, alternatively, this factor may be essential for viability or masked by genetic redundancy.

**Figure 5.**
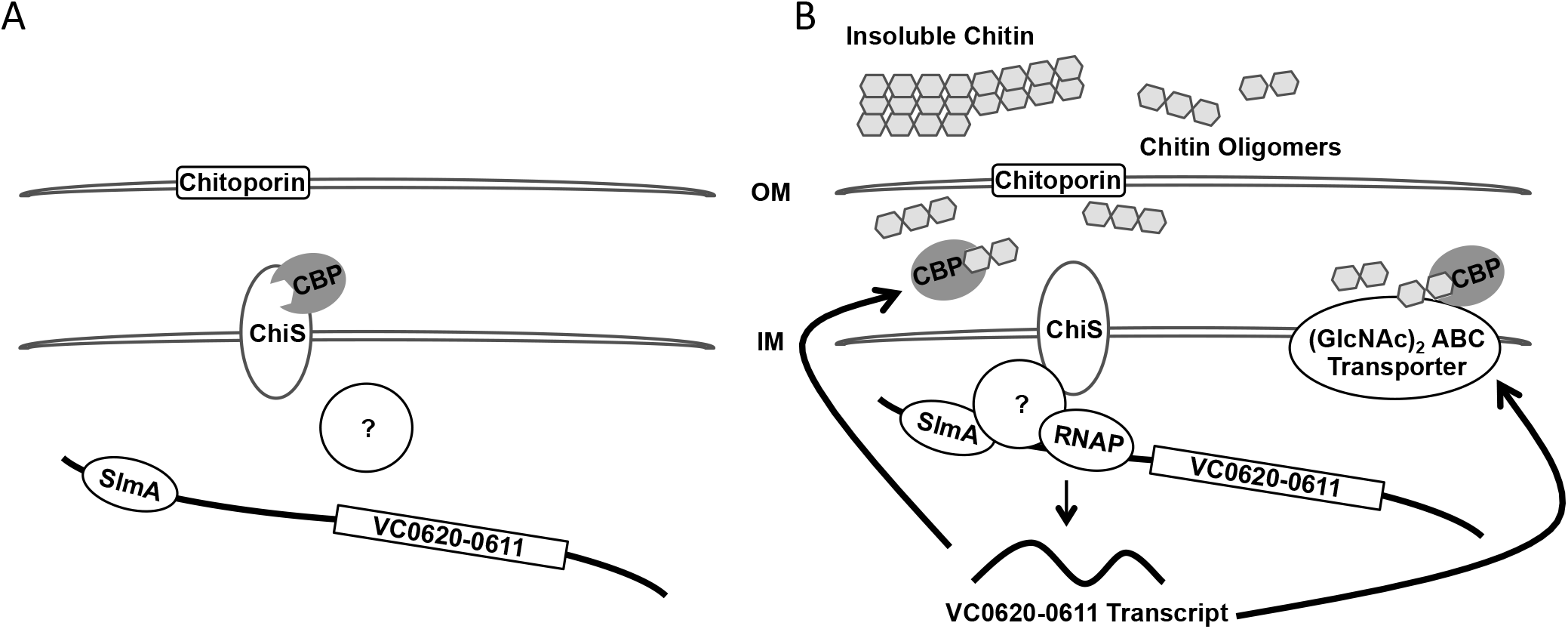
-Model for transcriptional activation of  *P_chb_*. (**A**) In the absence of chitin, CBP inhibits ChiS from activating expression of P_*chb*_. (**B**) In the presence of chitin, CBP binds chitin oligomers, which relieves repression of ChiS. We hypothesize that ChiS interacts with or activates another currently unidentified protein, which recruits RNA Polymerase to activate transcription of P_*chb*_ in a SlmA-dependent manner. SlmA may help to recruit the putative factor to the P_*chb*_ promoter or is an otherwise required coactivator of this locus. Upon expression of the *chb* operon, proteins involved in chitin uptake and utilization of chitobiose, including CBP and the chitobiose ABC permease, are synthesized.

Because ChiS is an orphan sensor kinase, the most logical coactivator would be a cognate response regulator. However, it has been shown that the conserved histidine and aspartate residues of ChiS that are critical for autophosphorylation and phosphorelay activity, respectively, are not important for activation of P_*chb*_ [20]. Thus, while the identity of the regulator that acts downstream of ChiS remains unknown, we believe it is unlikely to be a classical response regulator. It is possible that this coactivator bridges SlmA and RNA polymerase to mediate transcriptional activation (as depicted in **Fig. 5**). Alternatively, it is possible that the coactivator bends or structurally alters DNA in the *chb* promoter to enhance SlmA-dependent recruitment of RNA polymerase. Identifying and characterizing a putative coactivator for P_*chb*_ that acts downstream of ChiS will be the focus of future work.

Another possible mechanism of activation is that SlmA directly interacts with RNA polymerase through its ability to polymerize and spread on DNA [14]. We have two results, however, that diminish this as a likely model for activation. First, we show that moving the SBS as little as 3 bp in either direction within P_*chb*_ abolishes activation (**Fig. S7**). If DNA spreading were critical we would predict that this perturbation should have a limited effect on transcriptional activation. Further, we have shown that overexpression of SlmA can actually decrease P_*chb*_ transcription, which again would not be predicted by a DNA spreading model (**Fig. S10**).

We have shown that there are two transcriptional start sites for the *chb* locus. The short transcript is strongly induced in the presence of chitin oligosaccharides, while the long transcript is basally produced in the absence of chitin and is only modestly induced in its presence (2–4 fold). This indicates that there may be an additional level of regulation involving the 5’ untranslated region (UTR) of the *chb* transcript. A putative hairpin in P_*chb*_ also seemed to have some inhibitor effect on P_*chb*_ expression (**Fig. S5**). It is possible that the long transcript and hairpin play an important role in the basal expression of the *chb* operon, which is required for a rapid response to chitin oligosaccharides.

TetR proteins are generally transcriptional repressors, however, there are examples where these proteins can mediate transcriptional activation. One well-studied example is the LuxR protein in *Vibrio harveyi* (also known as HapR in *V. cholerae* and SmcR in *Vibrio vulnificus*) [21, 22]. SlmA is primarily characterized as a TetR family cell wall licensing factor and here we describe its additional role as a transcriptional activator. It has been hypothesized that transcriptional regulatory proteins arose from nucleoid-associated proteins [23]. Their role in binding DNA to properly structure the nucleoid may have evolved to aid in transcriptional regulation by contributing to the rearrangement of DNA structure at promoter regions [24]. One such example is integration host factor (IHF), which binds DNA and results in dramatic DNA bending (bend angle of ~120°), which can promote activation of regulated genes. SlmA binding results in subtle bending of DNA (~18°) [14]. Thus, SlmA may have evolved from a protein that structures the nucleoid into a cell division licensing factor as well as a transcriptional regulator. Additionally, since SlmA carries out both nucleoid occlusion and transcriptional activation, it is possible that these two activities affect one another. As a result, co-option of SlmA for regulation of P_*chb*_ may be a mechanism to integrate the cell division status of the cell with activation of chitin utilization.

To our knowledge, this is the first example of gene regulation by a nucleoid occlusion protein. A recent *in vitro* whole genome binding analysis identified 79 putative SBSs in *V. cholerae* [17]. We determined that ~25% (20/79) of these putative SBSs are in intergenic sites (including the one in P_*chb*_), while only ~12% of the genome constitutes intergenic sequence. By contrast, in *E. coli*, ~8.3% (2/24) of SBSs are in intergenic sites [12] and ~11% of the genome is intergenic. Also, analyzing binding sites for Noc, the nucleoid occlusion protein of *Bacillus subtilis*, ~12% (9/74) of binding sites are intergenic [25] and ~11% of the genome represents intergenic sequence. By contrast, for binding sites for the terminus macrodomain proteins MatP (known as *matS* sites) ~24% (6/25) are in intergenic regions [26]. Thus, it is possible that enriched binding of nucleoid-associated proteins at intergenic regions (as is the case for *matS* sites in *E. coli* and SBSs in *V. cholerae*) is because these sites are more flexible in regards to accommodating mutations compared to coding regions of the genome. Alternatively, SBSs may be enriched in intergenic sites in *V. cholerae* possibly to regulate the expression of additional genes. This will be the focus of future work. Preliminary RNA-seq analysis of a *slmA* mutant grown in rich medium, however, did not uncover any additional loci regulated by SlmA in *V. cholerae*. An effect of SlmA on gene regulation at P_*chb*_, however, was only uncovered by deletion of CBP or growth on chitin (both of which induce this locus). Thus, it may be that additional inducing cues are required to uncover a role for SlmA-dependent regulation at additional genetic loci. Nucleoid occlusion factors are present across diverse bacterial genera and it is tempting to speculate that these proteins may also participate in regulating gene expression in addition to their established roles in cell division licensing.

## MATERIALS AND METHODS

### Bacterial strains and culture conditions

*V. cholerae* strains were routinely grown in LB medium and on LB agar supplemented when necessary with Carbenicillin (20 or 50 μg/mL), Kanamycin (50 μg/mL), Spectinomycin (200 μg/mL), Trimethoprim (10 μg/mL), and/or Chloramphenicol (2 μg/mL). For growth on a defined carbon source, strains were grown in M9 minimal medium containing the indicated carbon source.

### Transposon mutagenesis

Transposon mutant libraries were generated with a Carb^R^ mini-Tn10 transposon exactly as previously described [27]. Briefly, the transposon mutagenesis plasmid pDL1086 was first mated into the P_*chb*_-*lacZ* or P_*chb*_-*lacZ* Δ*cbp* reporter strain. Transposition was induced by plating cultures on Carb containing media at 42°C. To screen colonies, plates also contained 40 μg/mL X-gal and 5 mM IPTG. IPTG was added to competitively inhibit the basal activity of the P_*chb*_-*lacZ* reporter. This made it easier to distinguish colonies where the locus was uninduced (e.g. P_*chb*_-lacZ) from when the locus was induced (e.g. P_*chb*_-*lacZ* Δ*cbp*).

### Generation of mutant strains and constructs

The parent strain used in this study was E7946 [28]. Mutant constructs were generated using splicing-by-overlap extension PCR exactly as previously described [29]. Transformations and cotransformations were carried out exactly as previously described [30]. Mutant strains were confirmed by PCR and/or sequencing. The SlmA expression plasmid was generated by cloning the WT SlmA gene (VC0214) into the NdeI and BamHI sites of a His expression vector (pHis-tev). Untagged SlmA was also cloned into the EcoRI and BamHI sites of the IPTG-inducible expression vector pMMB67EH. Site directed mutants of SlmA were subsequently generated from these constructs using parallel single primer reactions exactly as previously described [31]. All plasmid inserts were confirmed by sequencing. Error prone PCR of SlmA was carried out exactly as previously described [32]. A detailed description of mutant strains and primers used in this study are outlined in **Table S1** and **Table S2**, respectively.

### Measuring GFP fluorescence

GFP was measured exactly as previously described [33]. Briefly, strains were grown at 30°C overnight, washed, and resuspended in instant ocean medium to a final OD**600** of 1.0. Fluorescence was determined using a BioTek H1M plate reader with excitation set to 500 nm and emission at 540 nm.

### 5’ RACE

The 5’ end of transcripts were mapped using the SMARTer 5’/3’ RACE kit according to manufacturer’s instructions (Takara). The primers used are listed in **Table S2**.

### Growth Curves

Cells were grown in M9 minimal medium in the presence of 0.2% of the indicated carbon source at 37°C. Growth was kinetically monitored by measuring OD**600** on a BioTek H1M plate reader.

### Quantitative reverse transcriptase PCR (qRT-PCR)

RNA was isolated from cells using an RNEasy minikit according to manufacturer’s instructions (Qiagen). RNA was reverse transcribed using AffinityScript QPCR cDNA Synthesis Kit (Agilent). Quantitiative PCR was performed using iTaq Universal SYBR Green Supermix (Bio-Rad) with primers specific for the genes indicated (primers are listed in **Table S2**) and the reaction was monitored on a Step-One qPCR system.

### Electrophoretic Mobility Shift Assays (EMSAs)

EMSAs were performed essentially as previously described [29]. Briefly, probes were made by PCR. In the reaction we included Cy5-dCTP at a level that would result in incorporation of 1–2 Cy5 labeled nucleotides in the final probe. Binding reactions contained 20 mM Tris HCl pH 7.5, 100 mM KCl, 1 mM DTT, 5% glycerol, 0.1 mg/mL BSA, 0.1 mg/mL salmon sperm DNA, 2 nM Cy5 labeled DNA probe, and SlmA at the indicated concentrations in a 20 μL reaction volume. Reactions were incubated at room temperature for 30 minutes. Then, glycerol was added to a final concentration of 15% and 18 μL of each reaction was loaded onto a 5% polyacrylamide 0.5× TBE gel. The gel was run at 150V for 40 minutes in 0.5× TBE. Gels were imaged for Cy5 fluorescence on a Typhoon-9210 instrument or a BioRad ChemiDoc MP Imaging system.

### Western blot analysis

Cells were grown to mid-log in the presence of IPTG as an inducer where indicated. Cells were pelleted, resuspended and boiled in 1X SDS PAGE sample buffer and separated by polyacrylamide gel electrophoresis. Proteins were then transferred to PVDF and probed with rabbit polyconal α-FLAG (Sigma) or mouse monoclonal α-RpoA (Biolegend) primary antibodies. Blots were then incubated with α-rabbit or α-mouse secondary antibodies conjugated to IRdye 800CW as appropriate and imaged using an Odyssey classic LI-COR imaging system.

### Microscopy

Cells were grown to mid-log in LB medium and then mounted on 1% agarose pads. Cells were imaged using an Olympus IX83 phase contrast microscope.

## ACKNOWLEDGEMENTS

We would like to thank Dean Rowe-Magnus and Julia van Kessel for sharing protocols and reagents and for helpful discussions. This work was supported by US National Institutes of Health Grant AI118863 to A.B.D. and startup funds from the Indiana University College of Arts and Sciences.

## Supplementary information for

**Fig. S1.**
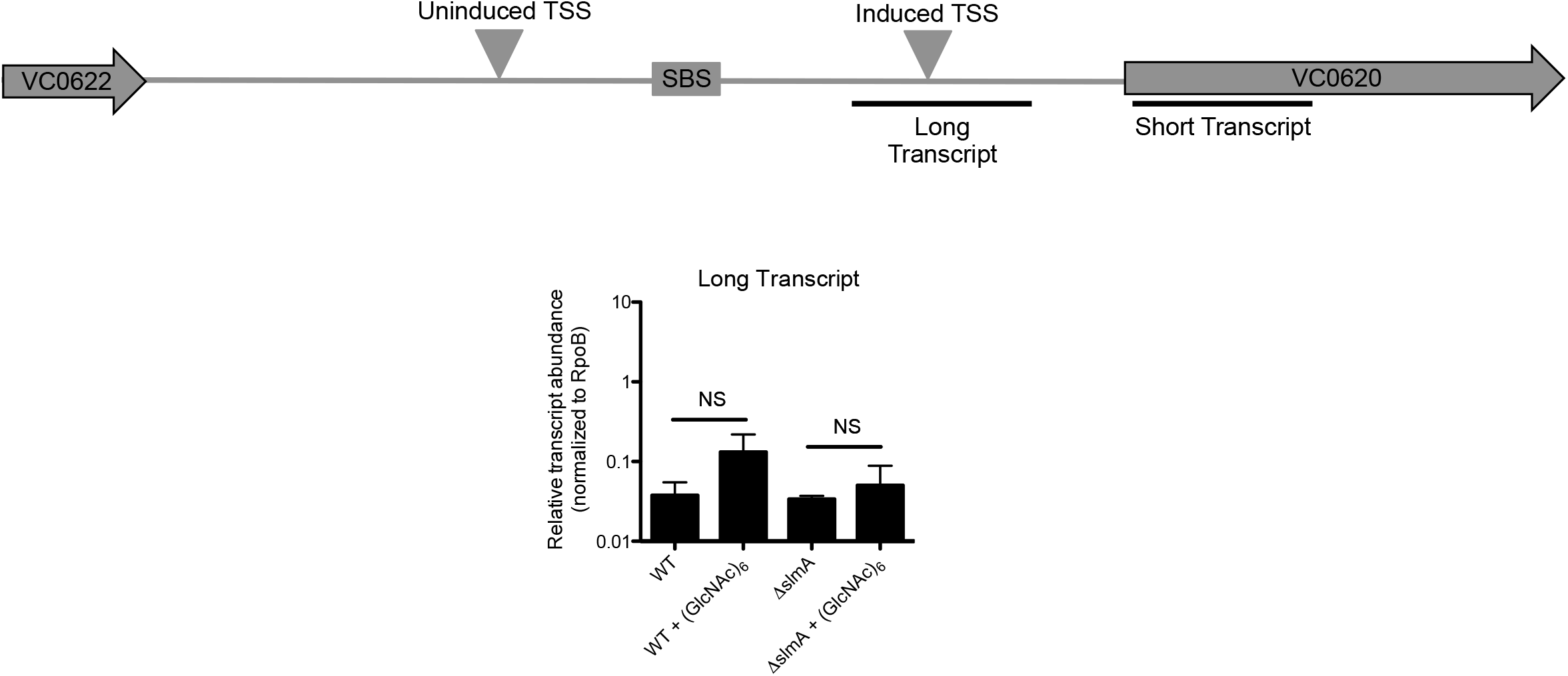
-Levels of the long “uninduced” *P_chb_* transcript do not change upon induction with (GlcNAc)_6_. Transcript levels were determined via qRT-PCR using primers specific for the long P_*chb*_ transcript. Data are shown as the mean ± SD and are from at least three independent biological replicates. **Fig. 1D** shows the qRT-PCR data for the short “induced” transcript.

**Fig. S2.**
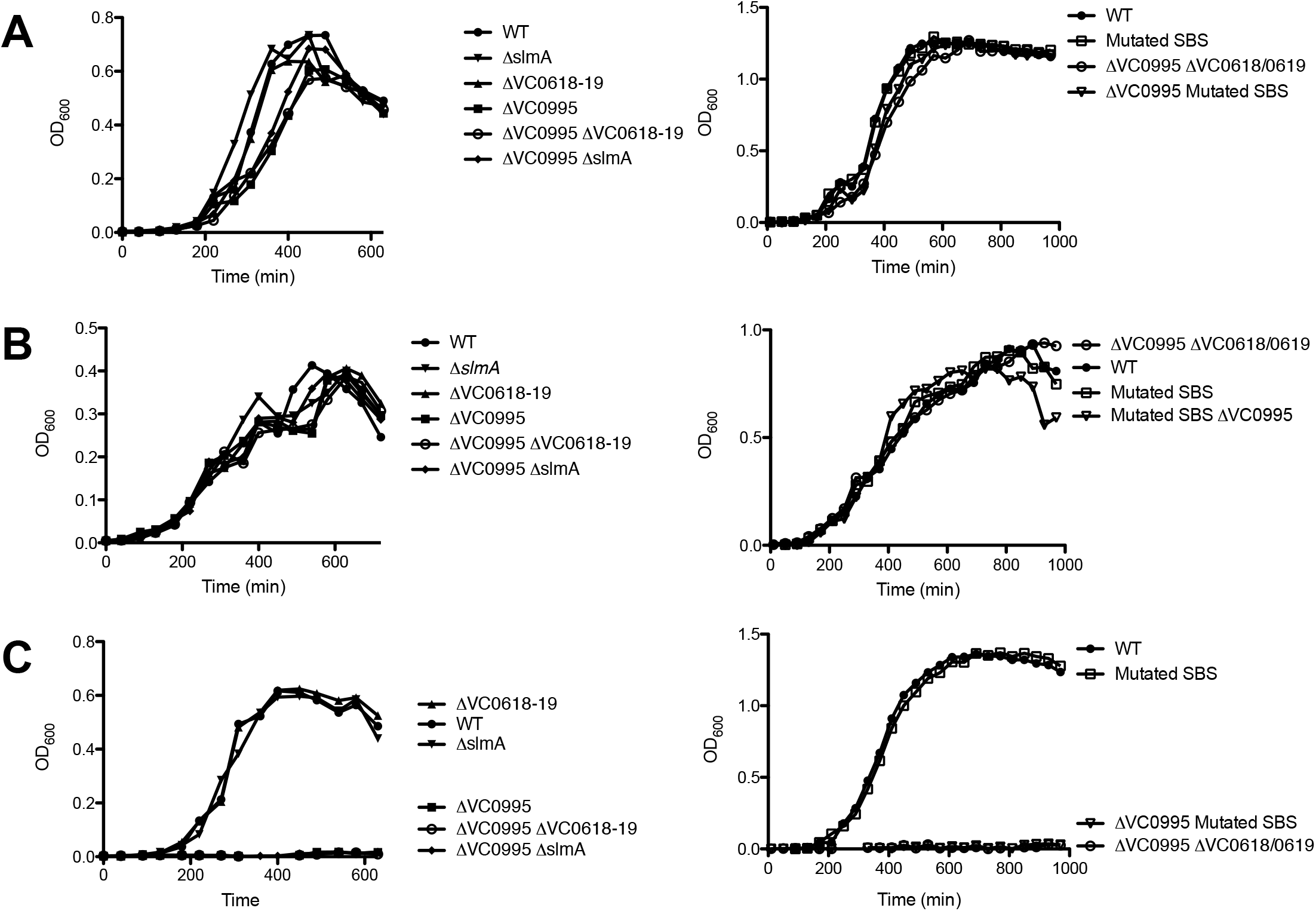
-*Mutant strains grow as expected on glucose, GlcNAc, and tryptone*. Growth curves of the indicated strains in M9 minimal medium containing 0.5% (**A**) glucose, (**B**) tryptone, or (**C**) *N*-acetylglucosamine (GlcNAc) as a sole carbon source.

**Fig. S3.**
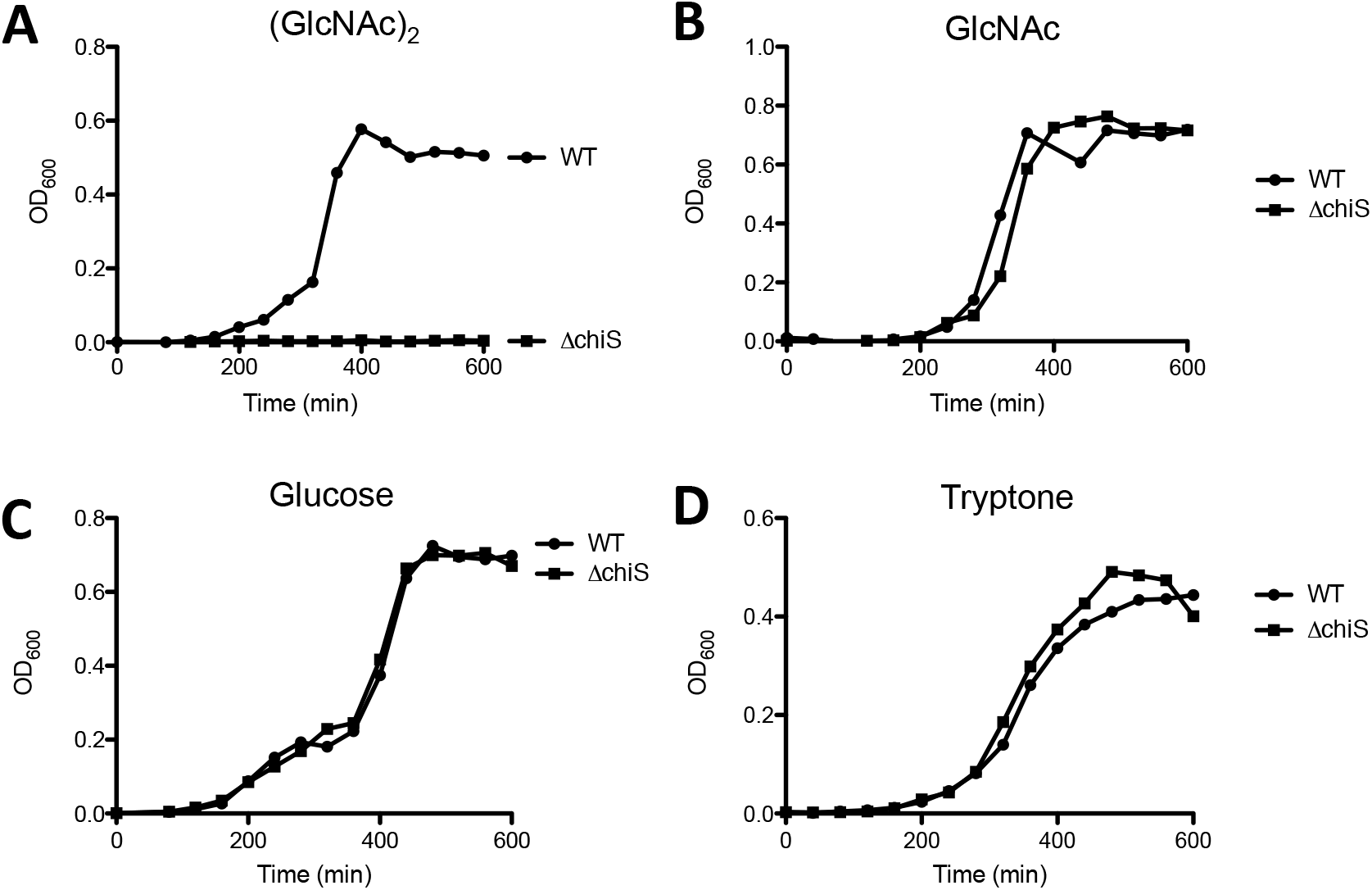
-ChiS is required for growth on chitobiose. Growth curves of the indicated strains in M9 minimal medium containing 0.5% (**A**) chiPobiose, (**B**) *N*-acetylglucosamine, (**C**) glucose, or (**D**) tryptone as a sole carbon source.

**Fig. S4.**
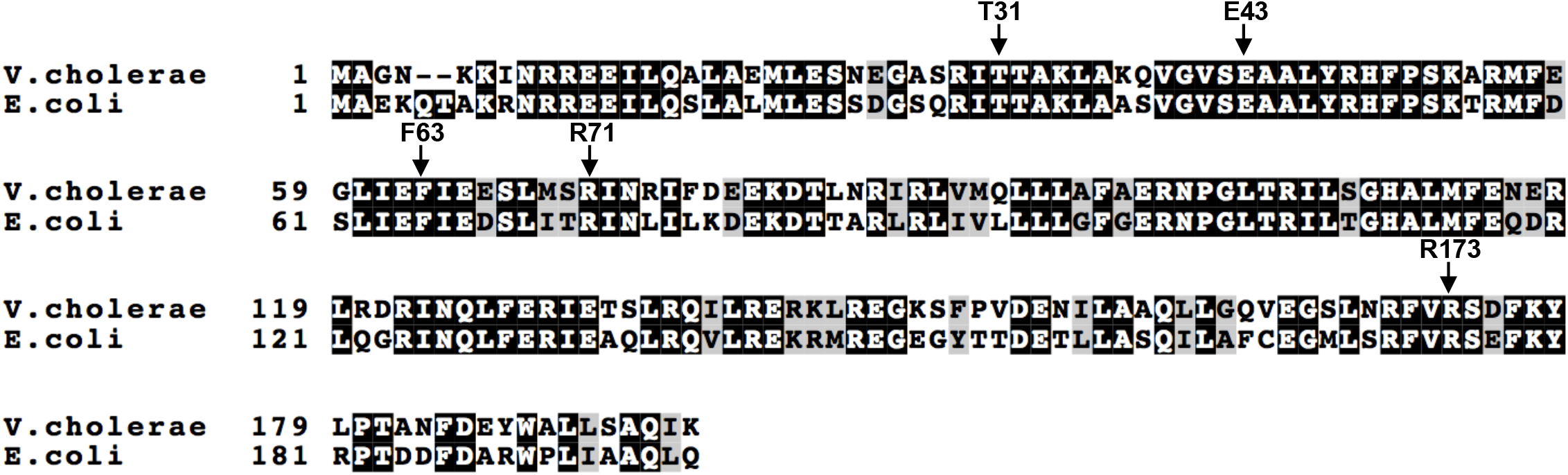
-*Sequence alignment of SlmA protein from V. cholerae and E. coli*. Residues highlighted in black are identical and those highlighted in gray are similar. Arrows indicate residues used in site-directed mutational analysis. T31 and E43 are involved in DNA-binding, F63 and R71 are involved in interaction with FtsZ, and R173 is involved in dimerization.

**Fig. S5.**
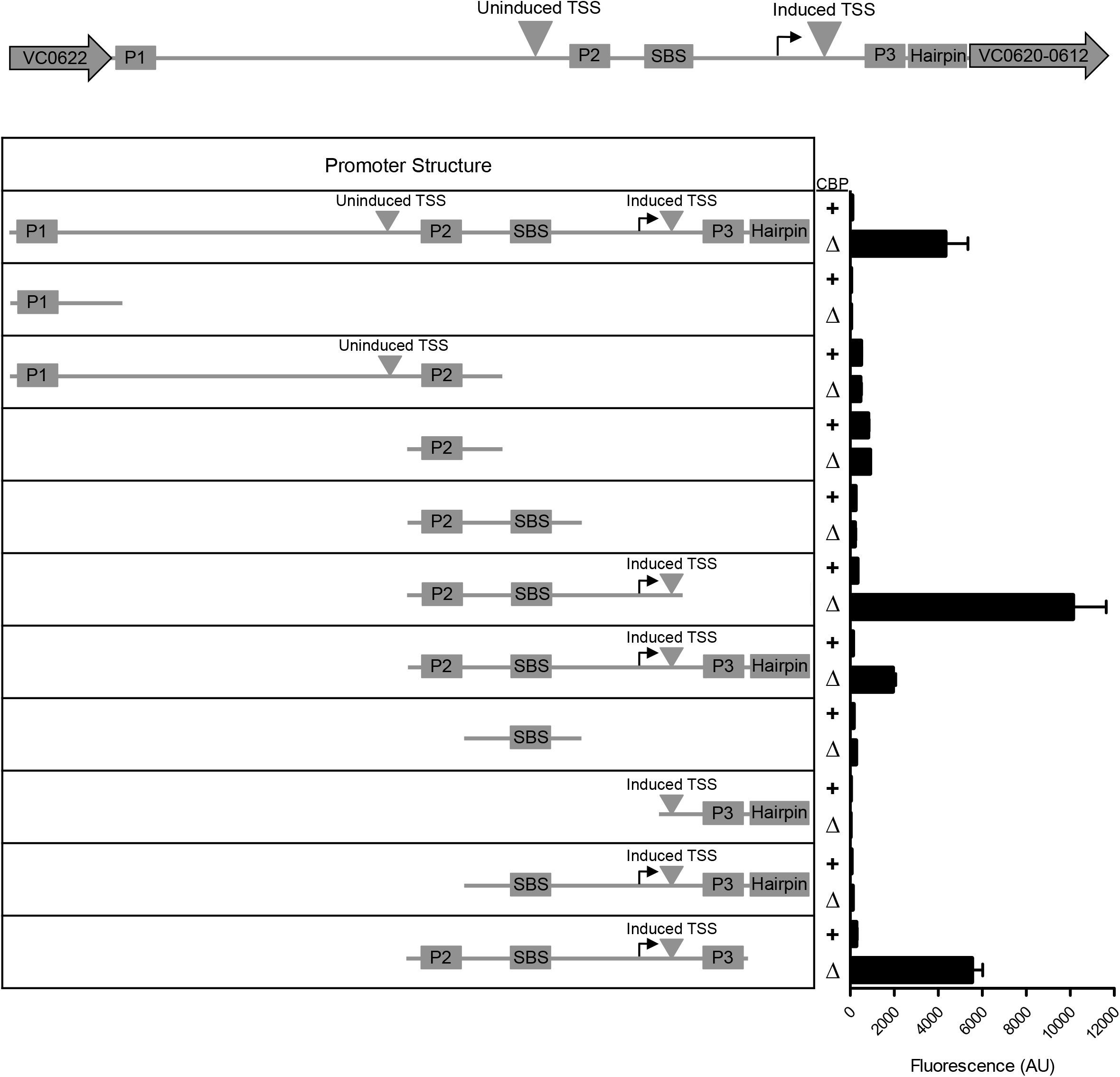
-Truncations narrow the region of *P_chb_* required for transcriptional activation. Promoters that were predicted computationally (Softberrry BPROM), are indicated as P1, P2, and P3, as well as a putative hairpin. All strains harbor the indicated region of *P_chb_* fused to *gfp* to serve as a transcriptional reporter. Also, strains are either intact for *cbp* (+) or harbor a *cbp* deletion (Δ). Data are shown as the mean ± SD and are from at least three independent biological replicates.

**Fig. S6.**
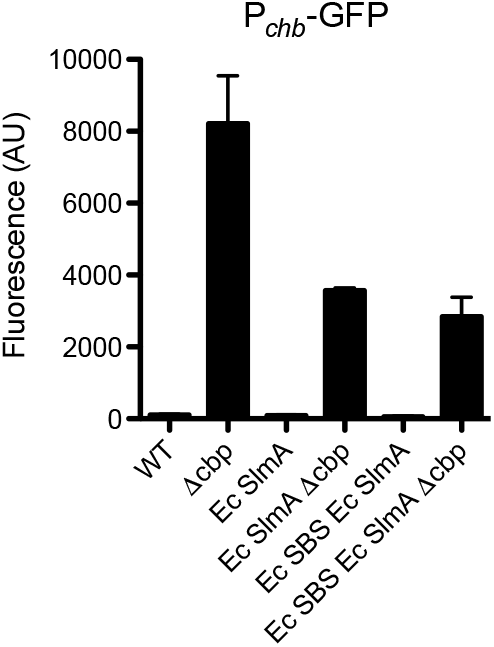
-An Ec SBS at *P_chb_* does not enhance activation of *P_chb_* by Ec SlmA. GFP fluorescence was measured in the indicated strains, all of which contain a P_*chb*_-*gfp* transcriptional reporter. “Ec SBS” indicates that the native SBS sequence in P_*chb*_ was swapped for the consensus SBS binding site for *Ec* SlmA. Data are from at least three independent biological replicates and shown as the mean ± SD. Please note that data from the first four bars is identical to that shown in **Fig. 1E**, and are included here to allow for easy comparison to the additional samples included in this figure.

**Fig. S7.**
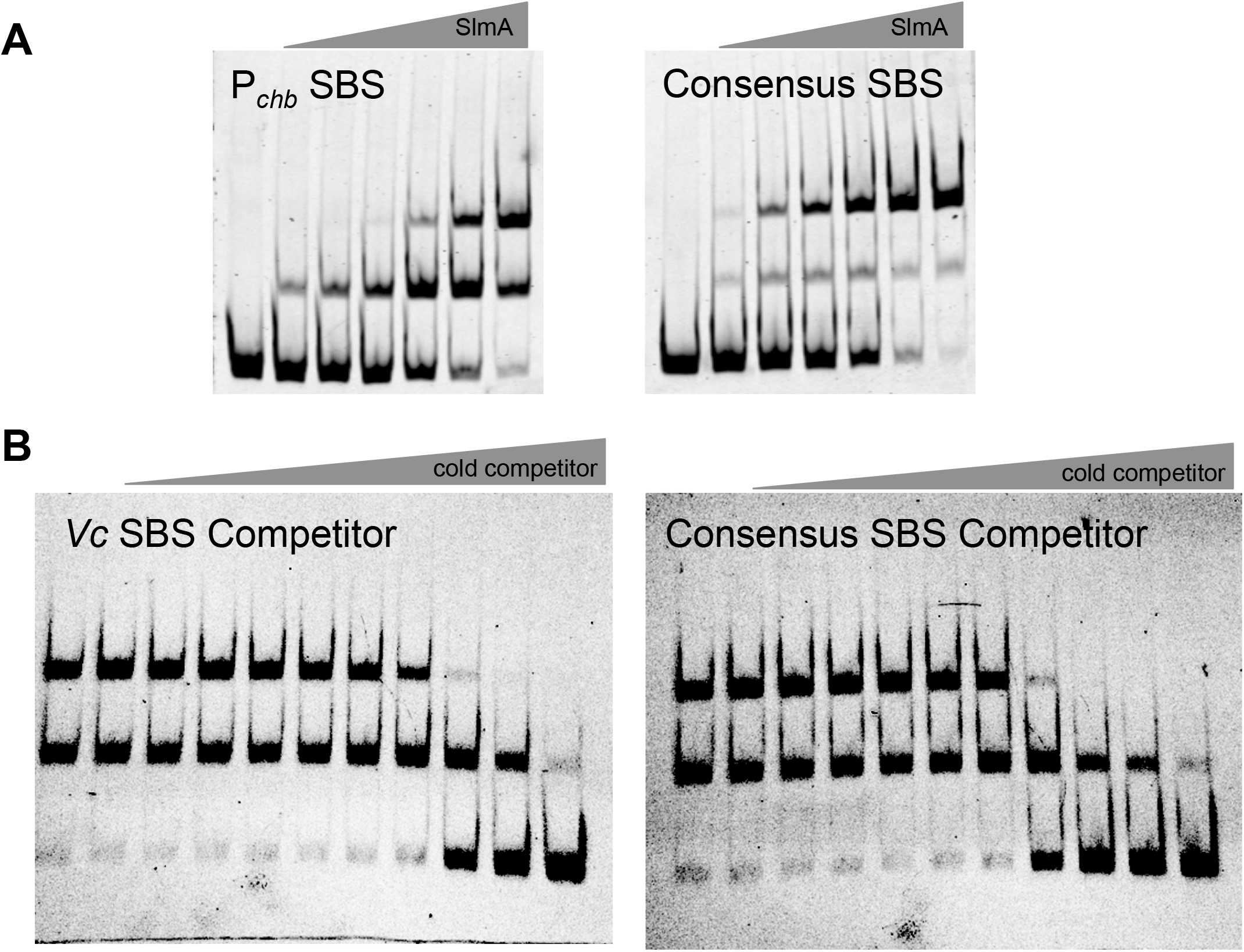
-SlmA has a higher affinity for a synthetic SBS that represents the consensus sequence for SlmA binding in E. coli compared to the SBS present in *P_chb_*. (**A**) EMSAs performed where the indicated SBS sequence containing probes were incubated with (from left to right): 0 nM, 7.5 nM, 15 nM, 30 nM, 60 nM, 120 nM, and 240 nM of purified SlmA. Data are representative of at least two independent experiments. (**B**) Labeled *Vc* SBS from *P_chb_* (0.2 nM) was incubated with increasing concentrations of unlabeled *Vc* SBS from P_*chb*_ or the consensus SBS from *E. coli*. Concentrations of unlabeled “cold competitor” probe from left to right are: 0 nM, 0.2 nM, 0.4 nM, 0.8 nM, 1.6 nM, 3.2 nM, 6.4 nM, 12.8 nM, 25.6 nM, 51.2 nM, and 200 nM.

**Fig. S8.**
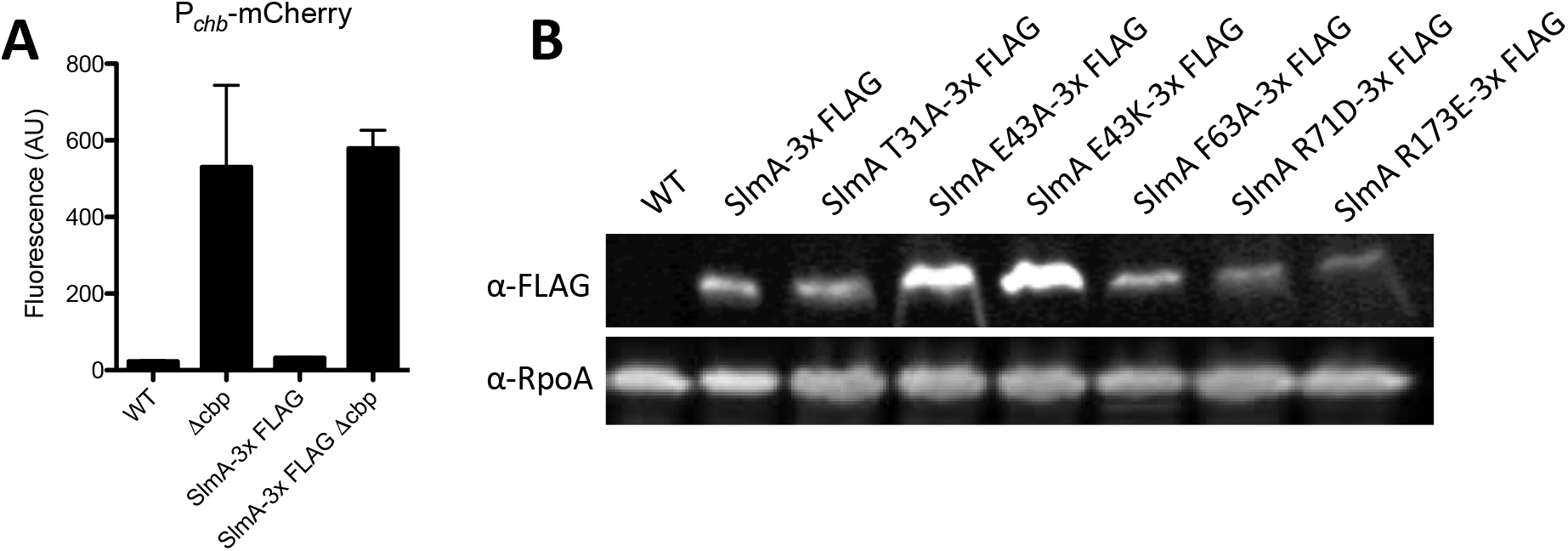
-Expression of SlmA mutant alleles is similar in vivo. (**A**) Fluorescence measurement of strains containing a P_*chb*_-mcherry reporter. Strains with SlmA-3x FLAG were at the native SlmA locus. Data indicates that the SlmA-3x FLAG construct is functional for P_*chb*_ activation. (**B**) *SlmA* site directed-mutants were engineered to contain a 3x FLAG tag at the native locus. Cell lysates of these strains were subjected to Western blot using a FLAG specific antibody and separately with an antibody to RpoA, which served as a loading control.

**Fig. S9.**
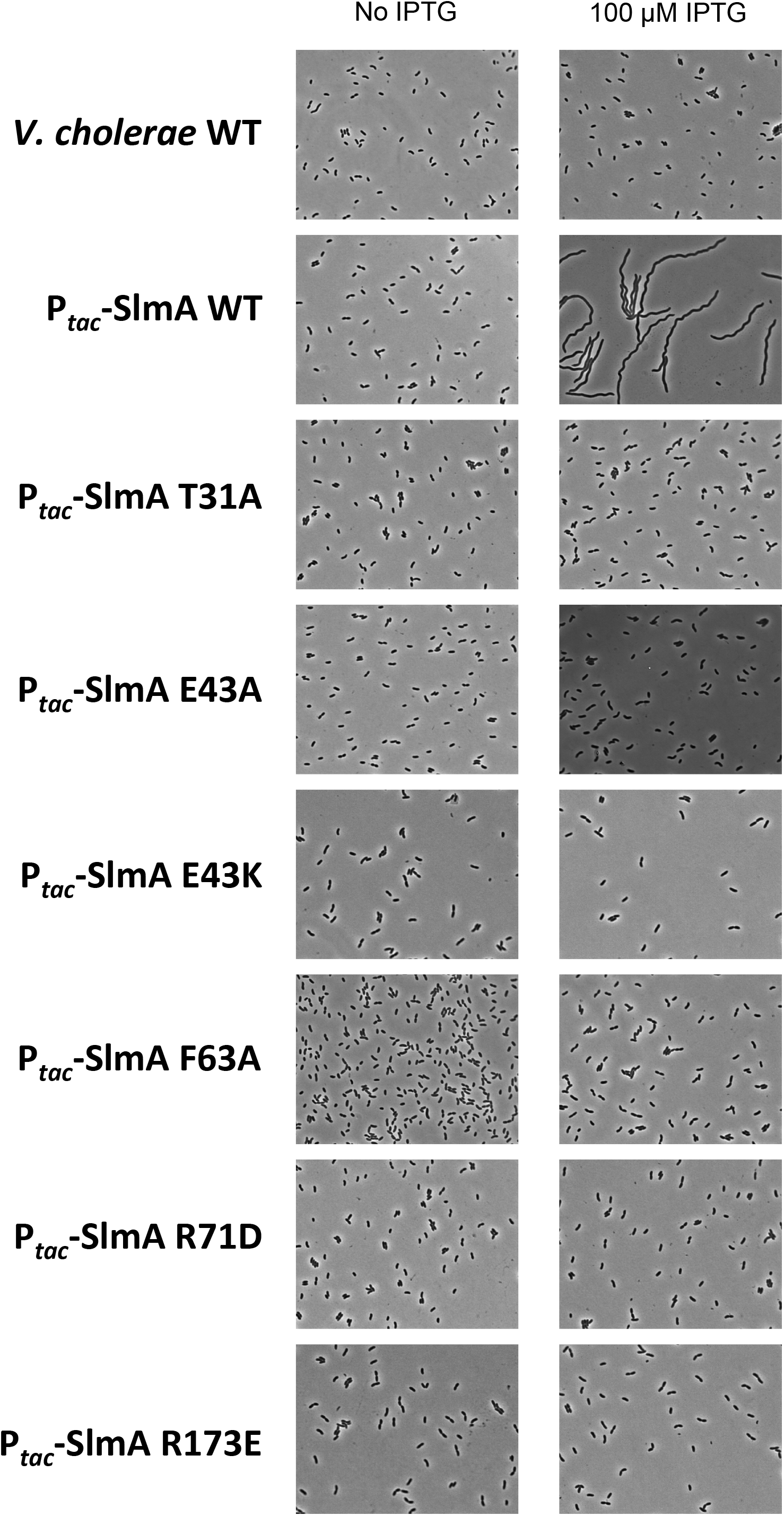
-Mutations in SlmA that reduce DNA binding or FtsZ interaction diminish nucleoid occlusion activity. Strains containing chromosomally integrated constructs for overexpression of the indicated SlmA mutants via an IPTG-inducible P_*tac*_ promoter were grown in the presence or absence of 100 μM IPTG and imaged by phase contrast microscopy. Data are representative of at least two independent experiments.

**Fig. S10.**
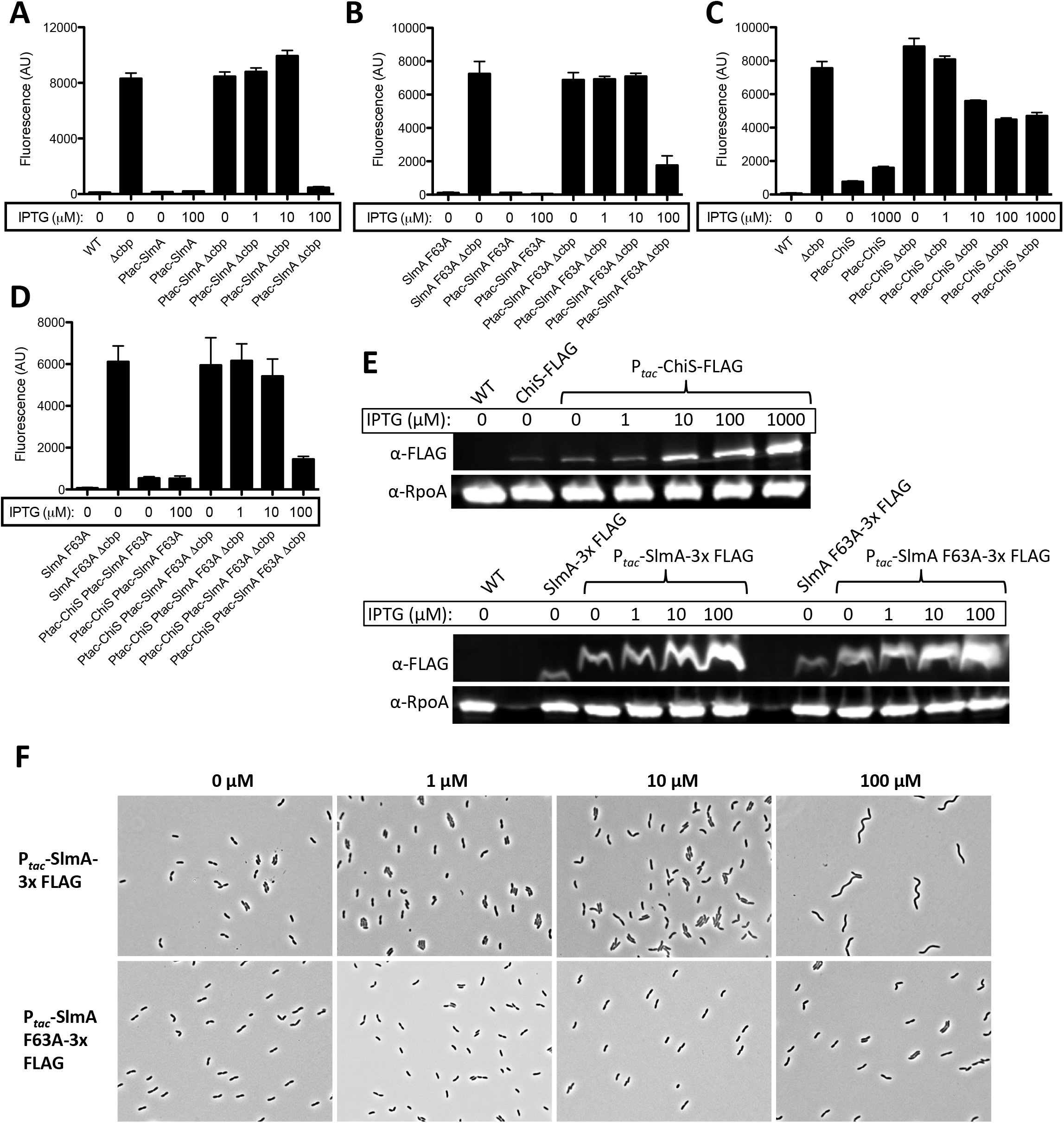
-Overexpression of SlmA and ChiS inhibits *P_chb_* activation. (**A-D**) GFP fluorescence was assessed in strains that harbor a P_*chb*_-*gfp* reporter as well as a chromosomally integrated construct for overexpression (P_*tac*_-X) of ChiS and/or SlmA as indicated. All strains were grown with the indicated concentration of IPTG. P_*tac*_-SlmA constructs contained a C-terminal triple FLAG tag, while P_*tac*_-ChiS constructs were untagged because the FLAG tag diminished the activity of this protein. Data are shown as the mean ± SD and are from at least three independent biological replicates. (**E**) Representative western blots of SlmA-3x FLAG and ChiS-FLAG overexpression using the same chromosomally integrated constructs used in **A-D** for SlmA and a ChiS-FLAG construct. (**F**) Representative phase contrast images to show the morphology of cells ectopically expressing WT SlmA or SlmA F63A using the same mutant constructs and concentrations of IPTG used in **A-D**.

**Fig. S11.**
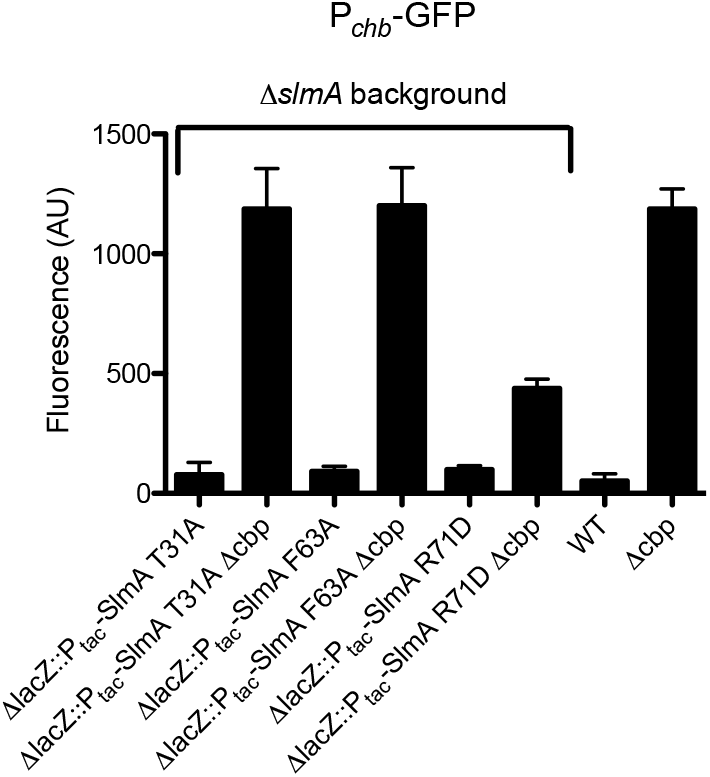
-Overexpression of SlmA is not sufficient to activate expression of *P_chb_*. GFP fluorescence was assessed in strains that harbor a P_*chb*_-*gfp* reporter as well as the other mutations indicated. All strains were grown with 10 μM IPTG to overexpress the indicated SlmA mutant proteins. Consistent with a model where SlmA requires another coactivator, overexpression of these SlmA variants was not sufficient to induce expression of P_*chb*_. Data are shown as the mean ± SD and are from at least three independent biological replicates.

**Fig. S12.**
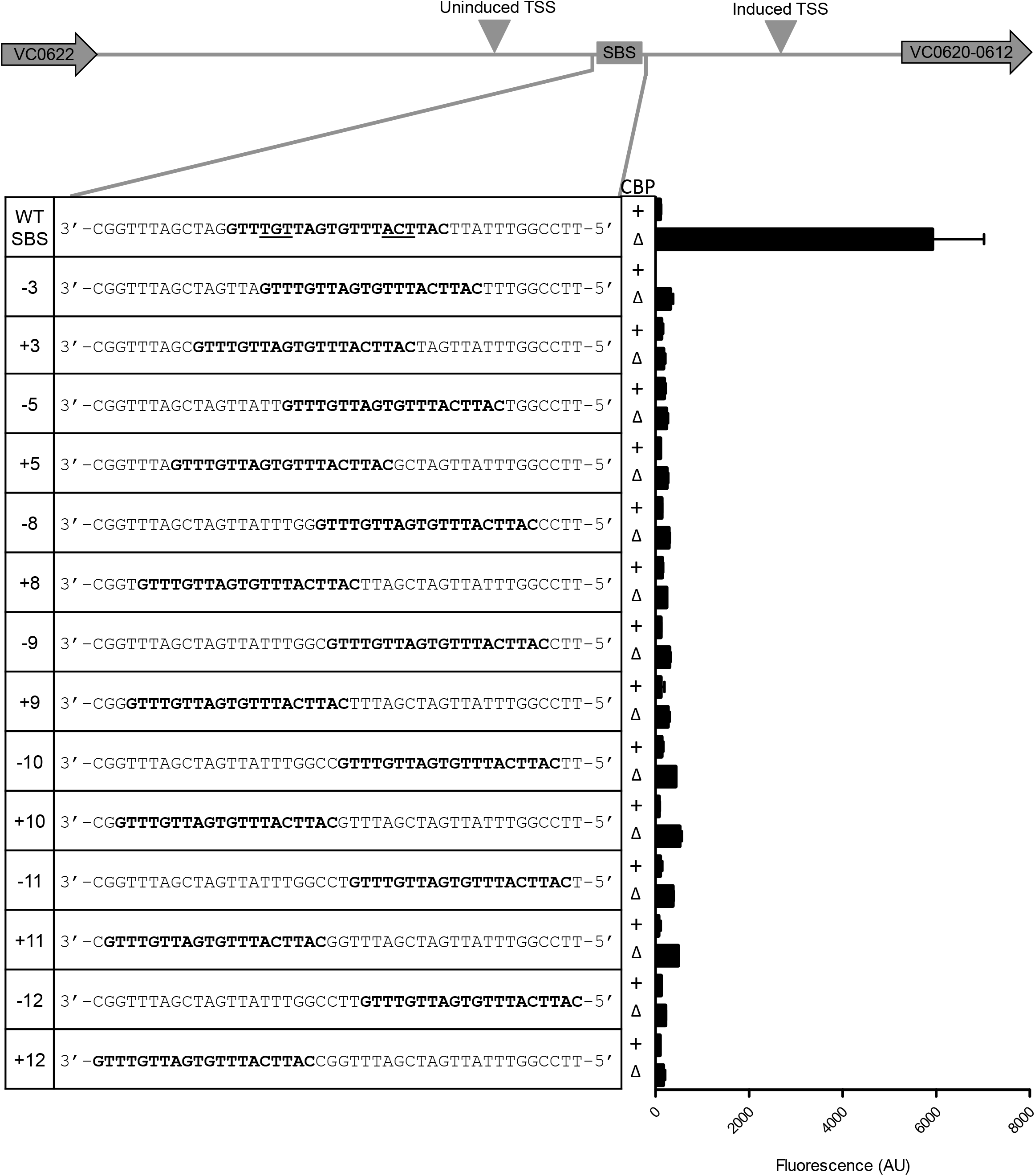
-The placement of the SBS in *P_chb_* is critical for transcriptional activation. All strains harbor a P_*chb*_-*gfp* reporter with the indicated mutation to move the SBS sequence as indicated. Each SBS mutation was tested in a background where *cbp* is intact (+) or deleted (Δ). Data are shown as the mean ± SD and are from at least three independent biological replicates.

**Fig. S13.**
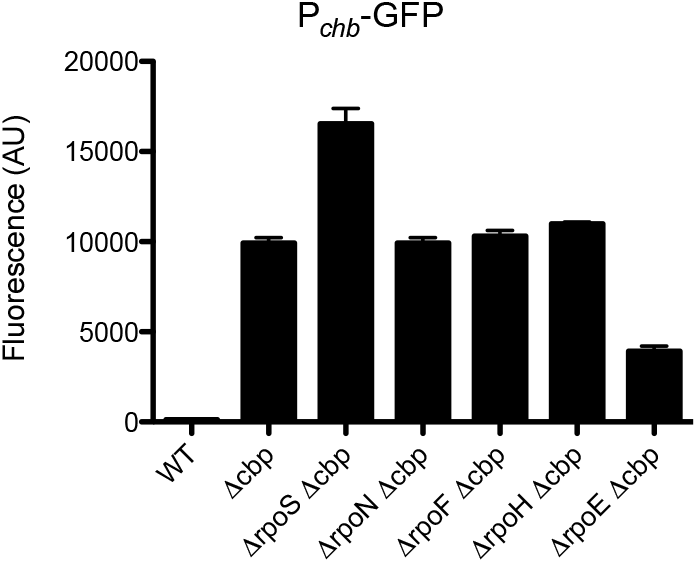
-Role of nonessential sigma factors in transcriptional activation of *P_chb_*. GFP fluorescence was assessed in strains that harbor a P_*chb*_-*gfp* reporter with the indicated sigma factor deleted. Data are shown as the mean ± SD and are from at least three independent biological replicates.

**Table S1.** -Strains used in this study

| Strain name in manuscript | Genotype and antibiotic resistances | Description | Reference / (strain#) |
| --- | --- | --- | --- |
| WT | E7946 Sm<sup>R</sup> | Wildtype *V. cholerae* O1 El Tor strain used throughout this study | (25)<br>(SAD030) |
| **Strains Used for Transposon Library** |  |  |  |
| Parent strain used for repressor transposon library | P<sub>chb</sub>-*lacZ*, Kan<sup>R</sup>; pDL1086 Amp<sup>R</sup> Cm<sup>R</sup> | CAK 062 parent harboring transposon vector pDL1086 | This Study (CAK 065 / SAD 1397) |
| Parent strain used for activator transposon library | P<sub>chb</sub>-*lacZ*, Kan<sup>R</sup>; ΔCBP::Spec<sup>R</sup>; pDL1086 Amp<sup>R</sup> Cm<sup>R</sup> | CAK 063 parent harboring transposon vector pDL1086 | This Study (CAK 066 / SAD 1398) |
| **Strains Used for Growth Curves** |  |  |  |
| ΔSlmA | Δ*slmA*::Kan<sup>R</sup> | Deletion of *slmA* replaced with Kan<sup>R</sup> | This Study (CAK 021 / SAD 1392) |
| ΔVC0995 | ΔVC1807::Spec<sup>R</sup>; ΔVC0995 | A deletion of VC0995 GlcNAc monosaccharide transporter and a deletion of VC1807 replaced with Spec<sup>R</sup> cassette | This Study (SAD 265) |
| ΔVC0995 ΔVC0618-19 | ΔVC1807::Spec<sup>R</sup>; ΔVC0995; ΔVC0618-19::Kan<sup>R</sup> | SAD 265 parent with a deletion of VC0618 and VC0619 replaced with Kan<sup>R</sup> | This Study (CAK 072 / SAD 1399) |
| ΔVC0995 Δ*slmA* | ΔVC1807::Spec<sup>R</sup>; ΔVC0995; Δ*slmA*::Tm<sup>R</sup> | SAD 265 parent with a deletion of *slmA* replaced with Tm<sup>R</sup> | This Study (CAK 073 / SAD 1400) |
| ΔVC0618-19 | ΔVC0618-19::Kan<sup>R</sup> | Deletion of VC0618 and VC0619 replaced with Kan<sup>R</sup> | This Study (SAD 382) |
| Scrambled SBS | P<sub>chb</sub> Scrambled SBS; ΔVC1807::Kan<sup>R</sup> | SBS at native P<sub>chb</sub> promoter was scrambled and VC1807 was replaced with Kan<sup>R</sup> | This Study (CAK 172 / SAD 1439) |
| Scrambled SBS ΔVC0995 | P<sub>chb</sub> Scrambled SBS; ΔVC1807::Kan<sup>R</sup>; ΔVC0995 | SAD 265 parent; SBS at native promoter was scrambled and Kan<sup>R</sup> cassette swapped for Spec<sup>R</sup> cassette at VC1807 | This Study (CAK 173 / SAD 1440) |
| Δ*chiS* | Δ*chiS*::Spec<sup>R</sup> | A deletion of *chiS* replaced with Spec<sup>R</sup> | This study (SAD 117) |
| **Strains Used for LacZ Activity** |  |  |  |
| Parent LacZ reporter | P<sub>chb</sub>-*lacZ*, Kan<sup>R</sup> | P<sub>chb</sub>-*lacZ* transcriptional reporter | This Study (CAK 062 / SAD 1395) |
| ΔCBP LacZ reporter | P<sub>chb</sub>-*lacZ*, Kan<sup>R</sup>; ΔCBP::Spec<sup>R</sup> | P<sub>chb</sub>-*lacZ* transcriptional reporter with a deletion of CBP replaced with Spec<sup>R</sup> cassette | This Study (CAK 063 / SAD 1396) |
| ΔCBP Δ*slmA* LacZ reporter | P<sub>chb</sub>-*lacZ*, Kan<sup>R</sup>; ΔCBP::Spec<sup>R</sup>; Δ*slmA*::Tm<sup>R</sup> | CAK 062 parent with a deletion of *slmA* replaced with Tm<sup>R</sup> and a deletion of CBP replaced with Spec<sup>R</sup> cassette | This Study (CAK 075 / SAD 1401) |
| Δ*ChiS* ΔCBP LacZ reporter | P<sub>chb</sub>-*lacZ*, Kan<sup>R</sup>; Δ*ChiS*::Tm<sup>R</sup>; ΔCBP::Spec<sup>R</sup> | CAK 062 parent with a deletion of *ChiS* replaced with Tm<sup>R</sup> and a deletion of CBP replaced with Spec<sup>R</sup> | This Study (CAK 209 / SAD 1460) |
| **Strains Used for P<sub>chb</sub>-GFP Assays** |  |  |  |
| WT GFP reporter, WT SBS GFP reporter | $\Delta lacZ::P_{chb}\text{-GFP Kan}^R$ | A deletion of *lacZ* replaced with $P_{chb}\text{-GFP}$ transcriptional fusion linked to $\text{Kan}^R$ | This Study (CAK 007 / SAD 590) |
| ΔCBP GFP reporter, WT SBS ΔCBP GFP reporter | $\Delta lacZ::P_{chb}\text{-GFP Kan}^R$; ΔCBP::Spec<sup>R</sup> | CAK 007 parent with a deletion of CBP replaced with Spec<sup>R</sup> cassette | This Study (CAK 009 / SAD 1390) |
| ΔCBP ΔChiS GFP reporter | $\Delta lacZ::P_{chb}\text{-GFP Kan}^R$; ΔCBP::Spec<sup>R</sup>; ΔChiS::Tm<sup>R</sup> | CAK 007 parent with a deletion of CBP replaced with Spec<sup>R</sup> cassette and a deletion of ChiS replaced with Tm<sup>R</sup> cassette | This Study (CAK 010 / SAD 1391) |
| ΔSlmA GFP reporter | $\Delta lacZ::P_{chb}\text{-GFP, Kan}^R$; $\Delta slmA::\text{Tm}^R$ | CAK 007 parent with a deletion of *slmA* replaced with Tm<sup>R</sup> cassette | This Study (CAK 024 / SAD 1393) |
| ΔSlmA ΔCBP GFP Reporter | $\Delta lacZ::P_{chb}\text{-GFP, Kan}^R$; $\Delta slmA::\text{Tm}^R$; ΔCBP::Spec<sup>R</sup> | CAK 007 parent with a deletion of *slmA* replaced with Tm<sup>R</sup> cassette and a deletion of CBP replaced with Spec<sup>R</sup> cassette | This Study (CAK 025 / 1394) |
| Scrambled SBS | $\Delta lacZ::P_{chb}\text{-GFP with scrambled SBS}$ | A deletion of *lacZ* replaced with $P_{chb}\text{-GFP}$ transcriptional fusion linked to $\text{Kan}^R$; SBS in promoter element was “scrambled” | This Study (CAK 088 / SAD 1402) |
| pMMB SlmA ΔCBP | $\Delta lacZ::P_{chb}\text{-GFP, Kan}^R$; $\Delta slmA::\text{Tm}^R$; ΔCBP::Spec<sup>R</sup>; pMMB67EH SlmA | CAK 025 parent harboring pMMB67EH with *slmA* insert | This Study (CAK 299 / SAD 1480) |
| pMMB ΔCBP | $\Delta lacZ::P_{chb}\text{-GFP, Kan}^R$; $\Delta slmA::\text{Tm}^R$; ΔCBP::Spec<sup>R</sup>; pMMB67EH empty vector | CAK 025 parent harboring pMMB67EH with no insert | This Study (CAK 300 / SAD 1481) |
| Scrambled SBS ΔCBP | $\Delta lacZ::P_{chb}\text{-GFP with scrambled SBS}$; ΔCBP::Spec<sup>R</sup> | A deletion of *lacZ* replaced with $P_{chb}\text{-GFP}$ transcriptional fusion linked to $\text{Kan}^R$; SBS in promoter element was “scrambled”; a deletion of CBP replaced with Spec<sup>R</sup> | This Study (CAK 089 / SAD 1403) |
| WT SlmA GFP reporter | $\Delta lacZ::P_{chb}\text{-GFP, Kan}^R$; $\Delta VC1807::\text{Spec}^R$ | CAK 024 parent with WT SlmA knocked back in at native locus; has the same phenotype as CAK 007 | This Study (CAK 093 / SAD 1404) |
| WT SlmA ΔCBP GFP reporter | $\Delta lacZ::P_{chb}\text{-GFP, Kan}^R$; ΔCBP::Spec<sup>R</sup> | CAK 024 parent with WT SlmA knocked back in at native locus and a deletion of CBP replaced with Spec<sup>R</sup>; has the same phenotype as CAK 009 | This Study (CAK 094 / SAD 1405) |
| SlmA T31A GFP reporter | $\Delta lacZ::P_{chb}\text{-GFP, Kan}^R$; $\Delta VC1807::\text{Spec}^R$; SlmA T31A | CAK 024 parent with SlmA T31A knocked back in at native locus | This Study (CAK 095 / SAD 1406) |
| SlmA T31A ΔCBP GFP reporter | $\Delta lacZ::P_{chb}\text{-GFP, Kan}^R$; ΔCBP::Spec<sup>R</sup>; SlmA T31A | CAK 024 parent with SlmA T31A knocked back in at native locus and a deletion of CBP replaced with Spec<sup>R</sup> | This Study (CAK 096 / SAD 1407) |
| SlmA E43A GFP reporter | $\Delta lacZ::P_{chb}\text{-GFP, Kan}^R$; $\Delta VC1807::\text{Spec}^R$; SlmA E43A | CAK 024 parent with SlmA E43A knocked back in at native locus | This Study (CAK 143 / SAD 1430) |
| SlmA E43A ΔCBP GFP reporter | $\Delta lacZ::P_{chb}$-GFP, Kan<sup>R</sup>; ΔCBP::Spec<sup>R</sup>; SlmA E43A | CAK 024 parent with SlmA E43A knocked back in at native locus and a deletion of CBP replaced with Spec<sup>R</sup> | This Study (CAK 144 / SAD 1431) |
| SlmA F63A GFP reporter | $\Delta lacZ::P_{chb}$-GFP, Kan<sup>R</sup>; ΔVC1807::Spec<sup>R</sup>; SlmA F63A | CAK 024 parent with SlmA F63A knocked back in at native locus | This Study (CAK 097 / SAD 1408) |
| SlmA F63A ΔCBP GFP reporter | $\Delta lacZ::P_{chb}$-GFP, Kan<sup>R</sup>; ΔCBP::Spec<sup>R</sup>; SlmA F63A | CAK 024 parent with SlmA F63A knocked back in at native locus and a deletion of CBP replaced with Spec<sup>R</sup> | This Study (CAK 098 / SAD 1409) |
| SlmA R71D GFP reporter | $\Delta lacZ::P_{chb}$-GFP, Kan<sup>R</sup>; ΔVC1807::Spec<sup>R</sup>; SlmA R71D | CAK 024 parent with SlmA R71D knocked back in at native locus | This Study (CAK 099 / SAD 1410) |
| SlmA R71D ΔCBP GFP reporter | $\Delta lacZ::P_{chb}$-GFP, Kan<sup>R</sup>; ΔCBP::Spec<sup>R</sup>; SlmA R71D | CAK 024 parent with SlmA R71D knocked back in at native locus and a deletion of CBP replaced with Spec<sup>R</sup> | This Study (CAK 108 / SAD 1413) |
| SlmA R173E GFP reporter | $\Delta lacZ::P_{chb}$-GFP, Kan<sup>R</sup>; ΔVC1807::Spec<sup>R</sup>; SlmA R173E | CAK 024 parent with SlmA R173E knocked back in at native locus | This Study (CAK 193 / SAD 1452) |
| SlmA R173E ΔCBP GFP reporter | $\Delta lacZ::P_{chb}$-GFP, Kan<sup>R</sup>; ΔCBP::Spec<sup>R</sup>; SlmA R173E | CAK 024 parent with SlmA R173E knocked back in at native locus and a deletion of CBP replaced with Spec<sup>R</sup> | This Study (CAK 194 / SAD 1453) |
| Ec SlmA GFP reporter | $\Delta lacZ::P_{chb}$-GFP, Kan<sup>R</sup>; ΔVC1807::Spec<sup>R</sup>; Ec SlmA | CAK 024 parent with SlmA from *E. coli* knocked back in at native locus | This Study (CAK 100 / SAD 1411) |
| Ec SlmA ΔCBP GFP reporter | $\Delta lacZ::P_{chb}$-GFP, Kan<sup>R</sup>; ΔCBP::Spec<sup>R</sup>; Ec SlmA | CAK 024 parent with SlmA from *E. coli* knocked back in at native locus and a deletion of CBP replaced with Spec<sup>R</sup> | This Study (CAK 101 / SAD 1412) |
| Ec SBS GFP reporter | $\Delta lacZ::P_{chb}$-GFP with Ec SBS | A deletion of *lacZ* replaced with *P<sub>chb</sub>*-GFP transcriptional fusion linked to Kan<sup>R</sup>; SBS in promoter element was replaced with *E. coli* consensus SBS | This Study (CAK 168 / SAD 1438) |
| Ec SBS GFP reporter ΔCBP | $\Delta lacZ::P_{chb}$-GFP with Ec SBS; ΔCBP::Spec<sup>R</sup> | CAK 168 parent with a deletion of CBP replaced with Spec<sup>R</sup> | This Study (CAK 197 / SAD 1455) |
| P1 | $\Delta lacZ::P1$ of *P<sub>chb</sub>*-GFP, Kan<sup>R</sup> | *P<sub>chb</sub>* truncation fused to GFP linked to Kan<sup>R</sup> | This Study (SAD 903) |
| P1 ΔCBP | $\Delta lacZ::P1$ of *P<sub>chb</sub>*-GFP, Kan<sup>R</sup>; ΔCBP::Spec<sup>R</sup> | *P<sub>chb</sub>* truncation fused to GFP linked to Kan<sup>R</sup> and a deletion of CBP replaced with Spec<sup>R</sup> | This Study (SAD 904) |
| P1+P2 | $\Delta lacZ::P1+P2$ of *P<sub>chb</sub>*-GFP, Kan<sup>R</sup> | *P<sub>chb</sub>* truncation fused to GFP linked to Kan<sup>R</sup> | This Study (SAD 912) |
| P1+P2 ΔCBP | $\Delta lacZ::P1+P2$ of *P<sub>chb</sub>*-GFP, Kan<sup>R</sup>; ΔCBP::Spec<sup>R</sup> | *P<sub>chb</sub>* truncation fused to GFP linked to Kan<sup>R</sup> and a deletion of CBP replaced with Spec<sup>R</sup> | This Study (SAD 913) |
| P2 only | $\Delta lacZ::P2$ only of *P<sub>chb</sub>*-GFP, Spec<sup>R</sup> | *P<sub>chb</sub>* truncation fused to GFP linked to Spec<sup>R</sup> | This Study (CAK 111 / SAD 1415) |
| P2 only ΔCBP | $\Delta lacZ::P_2$ only of $P_{chb^-}$-GFP; ΔCBP::Tm<sup>R</sup> | $P_{chb}$ truncation fused to GFP linked to Spec<sup>R</sup> and a deletion of CBP replaced with Tm<sup>R</sup> | This Study (CAK 112 / SAD 1416) |
| P2+SBS | $\Delta lacZ::P_2$+SBS of $P_{chb^-}$-GFP, Spec<sup>R</sup> | $P_{chb}$ truncation fused to GFP linked to Spec<sup>R</sup> | This Study (CAK 113 / SAD 1417) |
| P2+SBS ΔCBP | $\Delta lacZ::P_2$+SBS of $P_{chb^-}$-GFP, Spec<sup>R</sup>; ΔCBP::Tm<sup>R</sup> | $P_{chb}$ truncation fused to GFP linked to Spec<sup>R</sup> and a deletion of CBP replaced with Tm<sup>R</sup> | This Study (CAK 114 / SAD 1418) |
| P2+SBS more space | $\Delta lacZ::P_2$+SBS more space of $P_{chb^-}$-GFP, Spec<sup>R</sup> | $P_{chb}$ truncation fused to GFP linked to Spec<sup>R</sup> | This Study (CAK 116 / SAD 1419) |
| P2+SBS more space ΔCBP | $\Delta lacZ::P_2$+SBS more space of $P_{chb^-}$-GFP, Spec<sup>R</sup>; ΔCBP::Tm<sup>R</sup> | $P_{chb}$ truncation fused to GFP linked to Spec<sup>R</sup> and a deletion of CBP replaced with Tm<sup>R</sup> | This Study (CAK 117 / SAD 1420) |
| P2+P3 | $\Delta lacZ::P_2$+P3 of $P_{chb^-}$-GFP, Kan<sup>R</sup> | $P_{chb}$ truncation fused to GFP linked to Kan<sup>R</sup> | This Study (SAD 907) |
| P2+P3 ΔCBP | $\Delta lacZ::P_2$+P3 of $P_{chb^-}$-GFP, Kan<sup>R</sup>; ΔCBP::Spec<sup>R</sup> | $P_{chb}$ truncation fused to GFP linked to Kan<sup>R</sup> and a deletion of CBP replaced with Spec<sup>R</sup> | This Study (SAD 908) |
| P2+P3 internal region | $\Delta lacZ::P_2$+P3 internal region of $P_{chb^-}$-GFP, Spec<sup>R</sup> | $P_{chb}$ truncation fused to GFP linked to Spec<sup>R</sup> | This Study (CAK 129 / SAD 1425) |
| P2+P3 internal region ΔCBP | $\Delta lacZ::P_2$+P3 internal region of $P_{chb^-}$-GFP, Spec<sup>R</sup>; ΔCBP::Tm<sup>R</sup> | $P_{chb}$ truncation fused to GFP linked to Spec<sup>R</sup> and a deletion of CBP replaced with Tm<sup>R</sup> | This Study (CAK 130 / SAD 1426) |
| P3 | $\Delta lacZ::P_3$ of $P_{chb^-}$-GFP, Kan<sup>R</sup> | $P_{chb}$ truncation fused to GFP linked to Kan<sup>R</sup> | This Study (SAD 905) |
| P3 ΔCBP | $\Delta lacZ::P_3$ of $P_{chb^-}$-GFP, Kan<sup>R</sup>; ΔCBP::Spec<sup>R</sup> | $P_{chb}$ truncation fused to GFP linked to Kan<sup>R</sup> and a deletion of CBP replaced with Spec<sup>R</sup> | This Study (SAD 906) |
| P3 only more space | $\Delta lacZ::P_3$ only more space of $P_{chb^-}$-GFP, Spec<sup>R</sup> | $P_{chb}$ truncation fused to GFP linked to Spec<sup>R</sup> | This Study (CAK 119 / SAD 1421) |
| P3 only more space ΔCBP | $\Delta lacZ::P_3$ only more space of $P_{chb^-}$-GFP, Spec<sup>R</sup>; ΔCBP::Tm<sup>R</sup> | $P_{chb}$ truncation fused to GFP linked to Spec<sup>R</sup> and a deletion of CBP replaced with Tm<sup>R</sup> | This Study (CAK 120 / SAD 1422) |
| ΔHairpin | $\Delta lacZ::\Delta$hairpin of $P_{chb^-}$-GFP, Spec<sup>R</sup> | $P_{chb}$ truncation fused to GFP linked to Spec<sup>R</sup> | This Study (CAK 121 / SAD 1423) |
| ΔHairpin ΔCBP | $\Delta lacZ::\Delta$hairpin of $P_{chb^-}$-GFP, Spec<sup>R</sup>; ΔCBP::Tm<sup>R</sup> | $P_{chb}$ truncation fused to GFP linked to Spec<sup>R</sup> and a deletion of CBP replaced with Tm<sup>R</sup> | This Study (CAK 122 / SAD 1424) |
| -3 SBS | $\Delta lacZ::P_{chb^-}$-GFP, Kan<sup>R</sup>; -3 SBS | A deletion of *lacZ* replaced with $P_{chb^-}$-GFP transcriptional fusion linked to Kan<sup>R</sup>; SBS in promoter element was moved 3 bp downstream | This Study (CAK 179 / SAD 1441) |
| -3 SBS ΔCBP | $\Delta lacZ::P_{chb^-}$-GFP, Kan<sup>R</sup>; -3 SBS; ΔCBP::Spec<sup>R</sup> | A deletion of *lacZ* replaced with $P_{chb^-}$-GFP transcriptional fusion linked to Kan<sup>R</sup>; SBS in promoter element was moved 3 bp downstream; a deletion of CBP was replaced with Spec<sup>R</sup> | This Study (CAK 180 / SAD 1442) |
| +3 SBS | $\Delta lacZ::P_{chb}\text{-GFP, Kan}^R$; +3 SBS | A deletion of *lacZ* replaced with *P<sub>chb</sub>*-GFP transcriptional fusion linked to *Kan*<sup>R</sup>; SBS in promoter element was moved 3 bp upstream | This Study (CAK 183 / SAD 1445) |
| +3 SBS ΔCBP | $\Delta lacZ::P_{chb}\text{-GFP, Kan}^R$; +3 SBS; ΔCBP::*Spec*<sup>R</sup> | A deletion of *lacZ* replaced with *P<sub>chb</sub>*-GFP transcriptional fusion linked to *Kan*<sup>R</sup>; SBS in promoter element was moved 3 bp upstream; a deletion of CBP was replaced with *Spec*<sup>R</sup> | This Study (CAK 184 / SAD 1446) |
| -5 SBS | $\Delta lacZ::P_{chb}\text{-GFP, Kan}^R$; -5 SBS | A deletion of *lacZ* replaced with *P<sub>chb</sub>*-GFP transcriptional fusion linked to *Kan*<sup>R</sup>; SBS in promoter element was moved 5 bp downstream | This Study (CAK 181 / SAD 1443) |
| -5 SBS ΔCBP | $\Delta lacZ::P_{chb}\text{-GFP, Kan}^R$; -5 SBS; ΔCBP::*Spec*<sup>R</sup> | A deletion of *lacZ* replaced with *P<sub>chb</sub>*-GFP transcriptional fusion linked to *Kan*<sup>R</sup>; SBS in promoter element was moved 5 bp downstream; a deletion of CBP was replaced with *Spec*<sup>R</sup> | This Study (CAK 182 / SAD 1444) |
| +5 SBS | $\Delta lacZ::P_{chb}\text{-GFP, Kan}^R$; +5 SBS | A deletion of *lacZ* replaced with *P<sub>chb</sub>*-GFP transcriptional fusion linked to *Kan*<sup>R</sup>; SBS in promoter element was moved 5 bp upstream | This Study (CAK 185 / SAD 1447) |
| +5 SBS ΔCBP | $\Delta lacZ::P_{chb}\text{-GFP, Kan}^R$; +5 SBS; ΔCBP::*Spec*<sup>R</sup> | A deletion of *lacZ* replaced with *P<sub>chb</sub>*-GFP transcriptional fusion linked to *Kan*<sup>R</sup>; SBS in promoter element was moved 5 bp upstream; a deletion of CBP was replaced with *Spec*<sup>R</sup> | This Study (CAK 186 / SAD 1448) |
| -8 SBS | $\Delta lacZ::P_{chb}\text{-GFP, Kan}^R$; -8 SBS | A deletion of *lacZ* replaced with *P<sub>chb</sub>*-GFP transcriptional fusion linked to *Kan*<sup>R</sup>; SBS in promoter element was moved 8 bp downstream | This Study (CAK 217 / SAD 1463) |
| -8 SBS ΔCBP | $\Delta lacZ::P_{chb}\text{-GFP, Kan}^R$; -8 SBS; ΔCBP::*Spec*<sup>R</sup> | A deletion of *lacZ* replaced with *P<sub>chb</sub>*-GFP transcriptional fusion linked to *Kan*<sup>R</sup>; SBS in promoter element was moved 8 bp downstream; a deletion of CBP replaced with *Spec*<sup>R</sup> | This Study (CAK 225 / SAD 1471) |
| +8 SBS | $\Delta lacZ::P_{chb}\text{-GFP, Kan}^R$; +8 SBS | A deletion of *lacZ* replaced with *P<sub>chb</sub>*-GFP transcriptional fusion linked to *Kan*<sup>R</sup>; SBS in promoter element was moved 8 bp upstream | This Study (CAK 216 / SAD 1462) |
| +8 SBS ΔCBP | $\Delta lacZ::P_{chb}\text{-GFP, Kan}^R$; +8 SBS; ΔCBP::*Spec*<sup>R</sup> | A deletion of *lacZ* replaced with *P<sub>chb</sub>*-GFP transcriptional fusion linked to *Kan*<sup>R</sup>; SBS in promoter element was moved 8 bp upstream; a deletion of CBP replaced with *Spec*<sup>R</sup> | This Study (CAK 224 / SAD 1470) |
| -9 SBS | $\Delta lacZ::P_{chb}\text{-GFP, Kan}^R$; -9 SBS | A deletion of *lacZ* replaced with *P<sub>chb</sub>* -GFP transcriptional fusion linked to *Kan* <sup>R</sup>; SBS in promoter element was moved 9 bp downstream | This Study (CAK 219 / SAD 1465) |
| -9 SBS ΔCBP | $\Delta lacZ::P_{chb}$-GFP, Kan<sup>R</sup>; -9 SBS; ΔCBP::Spec<sup>R</sup> | A deletion of *lacZ* replaced with $P_{chb}$-GFP transcriptional fusion linked to Kan<sup>R</sup>; SBS in promoter element was moved 9 bp downstream; a deletion of CBP replaced with Spec<sup>R</sup> | This Study (CAK 227 / SAD 1473) |
| +9 SBS | $\Delta lacZ::P_{chb}$-GFP, Kan<sup>R</sup>; +9 SBS | A deletion of *lacZ* replaced with $P_{chb}$-GFP transcriptional fusion linked to Kan<sup>R</sup>; SBS in promoter element was moved 9 bp upstream | This Study (CAK 218 / SAD 1464) |
| +9 SBS ΔCBP | $\Delta lacZ::P_{chb}$-GFP, Kan<sup>R</sup>; +9 SBS; ΔCBP::Spec<sup>R</sup> | A deletion of *lacZ* replaced with $P_{chb}$-GFP transcriptional fusion linked to Kan<sup>R</sup>; SBS in promoter element was moved 9 bp upstream; a deletion of CBP replaced with Spec<sup>R</sup> | This Study (CAK 226 / SAD 1472) |
| -10 SBS | $\Delta lacZ::P_{chb}$-GFP, Kan<sup>R</sup>; -10 SBS | A deletion of *lacZ* replaced with $P_{chb}$-GFP transcriptional fusion linked to Kan<sup>R</sup>; SBS in promoter element was moved 10 bp downstream | This Study (CAK 200 / SAD 1456) |
| -10 SBS ΔCBP | $\Delta lacZ::P_{chb}$-GFP, Kan<sup>R</sup>; -10 SBS; ΔCBP::Spec<sup>R</sup> | A deletion of *lacZ* replaced with $P_{chb}$-GFP transcriptional fusion linked to Kan<sup>R</sup>; SBS in promoter element was moved 10 bp downstream; a deletion of CBP was replaced with Spec<sup>R</sup> | This Study (CAK 201 / SAD 1457) |
| +10 SBS | $\Delta lacZ::P_{chb}$-GFP, Kan<sup>R</sup>; +10 SBS | A deletion of *lacZ* replaced with $P_{chb}$-GFP transcriptional fusion linked to Kan<sup>R</sup>; SBS in promoter element was moved 10 bp upstream | This Study (CAK 202 / SAD 1458) |
| +10 SBS ΔCBP | $\Delta lacZ::P_{chb}$-GFP, Kan<sup>R</sup>; +10 SBS; ΔCBP::Spec<sup>R</sup> | A deletion of *lacZ* replaced with $P_{chb}$-GFP transcriptional fusion linked to Kan<sup>R</sup>; SBS in promoter element was moved 10 bp upstream; a deletion of CBP was replaced with Spec<sup>R</sup> | This Study (CAK 203 / SAD 1459) |
| -11 SBS | $\Delta lacZ::P_{chb}$-GFP, Kan<sup>R</sup>; -11 SBS | A deletion of *lacZ* replaced with $P_{chb}$-GFP transcriptional fusion linked to Kan<sup>R</sup>; SBS in promoter element was moved 11 bp downstream | This Study (CAK 221 / SAD 1467) |
| -11 SBS ΔCBP | $\Delta lacZ::P_{chb}$-GFP, Kan<sup>R</sup>; -11 SBS; ΔCBP::Spec<sup>R</sup> | A deletion of *lacZ* replaced with $P_{chb}$-GFP transcriptional fusion linked to Kan<sup>R</sup>; SBS in promoter element was moved 11 bp downstream; a deletion of CBP replaced with Spec<sup>R</sup> | This Study (CAK 229 / SAD 1475) |
| +11 SBS | $\Delta lacZ::P_{chb}$-GFP, Kan<sup>R</sup>; +11 SBS | A deletion of *lacZ* replaced with $P_{chb}$ -GFP transcriptional fusion linked to Kan<sup>R</sup>; SBS in promoter element was moved 11 bp upstream | This Study (CAK 220 / SAD 1466) |
| +11 SBS ΔCBP | $\Delta lacZ::P_{chb}\text{-GFP, Kan}^R$; +11 SBS; ΔCBP::Spec<sup>R</sup> | A deletion of *lacZ* replaced with *P<sub>chb</sub>*-GFP transcriptional fusion linked to Kan<sup>R</sup>; SBS in promoter element was moved 11 bp upstream; a deletion of CBP replaced with Spec<sup>R</sup> | This Study (CAK 228 / SAD 1474) |
| -12 SBS | $\Delta lacZ::P_{chb}\text{-GFP, Kan}^R$; -12 SBS | A deletion of *lacZ* replaced with *P<sub>chb</sub>*-GFP transcriptional fusion linked to Kan<sup>R</sup>; SBS in promoter element was moved 12 bp downstream | This Study (CAK 223 / SAD 1469) |
| -12 SBS ΔCBP | $\Delta lacZ::P_{chb}\text{-GFP, Kan}^R$; -12 SBS; ΔCBP::Spec<sup>R</sup> | A deletion of *lacZ* replaced with *P<sub>chb</sub>*-GFP transcriptional fusion linked to Kan<sup>R</sup>; SBS in promoter element was moved 12 bp downstream; a deletion of CBP replaced with Spec<sup>R</sup> | This Study (CAK 231 / SAD 1477) |
| +12 SBS | $\Delta lacZ::P_{chb}\text{-GFP, Kan}^R$; +12 SBS | A deletion of *lacZ* replaced with *P<sub>chb</sub>*-GFP transcriptional fusion linked to Kan<sup>R</sup>; SBS in promoter element was moved 12 bp upstream | This Study (CAK 222 / SAD 1468) |
| +12 SBS ΔCBP | $\Delta lacZ::P_{chb}\text{-GFP, Kan}^R$; +12 SBS; ΔCBP::Spec<sup>R</sup> | A deletion of *lacZ* replaced with *P<sub>chb</sub>*-GFP transcriptional fusion linked to Kan<sup>R</sup>; SBS in promoter element was moved 12 bp upstream; a deletion of CBP replaced with Spec<sup>R</sup> | This Study (CAK 230 / SAD 1476) |
| $\Delta lacZ::P_{tac}\text{-slmA}$ T31A Δ*slmA* | $\Delta lacZ::P_{tac}\text{-SlmA T31A Spec}^R$, Δ*slmA*::Tm<sup>R</sup>, ΔVC1807::P<sub>chb</sub>-gfp Kan<sup>R</sup> | Overexpression construct for the indicated SlmA allele was integrated at the *lacZ* locus, while the *P<sub>chb</sub>* reporter was integrated at the VC1807 locus, strain also has a deletion of *slmA* | This study (SAD1239) |
| $\Delta lacZ::P_{tac}\text{-slmA}$ T31A Δ*cbp* Δ*slmA* | $\Delta lacZ::P_{tac}\text{-SlmA T31A Spec}^R$, Δ*slmA*::Tm<sup>R</sup>, Δ*cbp*::Carb<sup>R</sup>, ΔVC1807::P<sub>chb</sub>-gfp | Overexpression construct for the indicated SlmA allele was integrated at the *lacZ* locus, while the *P<sub>chb</sub>* reporter was integrated at the VC1807 locus, strain also has a deletion of *slmA* and *cbp* | This study (SAD1242) |
| $\Delta lacZ::P_{tac}\text{-slmA}$ F63A Δ*slmA* | $\Delta lacZ::P_{tac}\text{-SlmA F63A, ΔslmA::Tm}^R$, ΔVC1807::P<sub>chb</sub>-gfp | Overexpression construct for the indicated SlmA allele was integrated at the *lacZ* locus, while the *P<sub>chb</sub>* reporter was integrated at the VC1807 locus, strain also has a deletion of *slmA* | This study (SAD1240) |
| $\Delta lacZ::P_{tac}\text{-slmA}$ F63A Δ*cbp* Δ*slmA* | $\Delta lacZ::P_{tac}\text{-SlmA F63A, ΔslmA::Tm}^R$, Δ*cbp*::Carb<sup>R</sup>, ΔVC1807::P<sub>chb</sub>-gfp | Overexpression construct for the indicated SlmA allele was integrated at the *lacZ* locus, while the *P<sub>chb</sub>* reporter was integrated at the VC1807 locus, strain also has a deletion of *slmA* and *cbp* | This study (SAD1243) |
| $\Delta lacZ::P_{tac}-slmA$<br>R71D $\Delta slmA$ | $\Delta lacZ::P_{tac}-SlmA$ R71D,<br>$\Delta slmA::Tm^R$,<br>$\Delta VC1807::P_{chb}-gfp$ | Overexpression construct for the indicated SlmA allele was integrated at the lacZ locus, while the $P_{chb}$ reporter was integrated at the VC1807 locus, strain also has a deletion of *slmA* | This study<br>(SAD1241) |
| $\Delta lacZ::P_{tac}-slmA$<br>R71D $\Delta cbp \Delta slmA$ | $\Delta lacZ::P_{tac}-SlmA$ R71D,<br>$\Delta slmA::Tm^R$, $\Delta cbp::Carb^R$,<br>$\Delta VC1807::P_{chb}-gfp$ | Overexpression construct for the indicated SlmA allele was integrated at the lacZ locus, while the $P_{chb}$ reporter was integrated at the VC1807 locus, strain also has a deletion of *slmA* and *cbp* | This study<br>(SAD1244) |
| $P_{chb}$ 112 bp *lacZ*<br>swap-gfp | $\Delta lacZ::P_{chb}-gfp$ 112 bp<br>swap Kan<sup>R</sup> | $\Delta lacZ::P_{chb}-gfp$ with nucleotides 612-724 swapped with an intergenic region from the *lacZ* gene linked to Kan<sup>R</sup> | This Study<br>(CAK 337 /<br>SAD 1482) |
| $P_{chb}$ 112 bp *lacZ*<br>swap-gfp $\Delta cbp$ | $\Delta lacZ::P_{chb}-gfp$ 112 bp<br>swap Kan<sup>R</sup>, $\Delta CBP::Spec^R$ | $\Delta lacZ::P_{chb}-gfp$ with nucleotides 612-724 swapped with an intergenic region from the *lacZ* gene linked to Kan<sup>R</sup>; deletion of CBP replaced with Spec<sup>R</sup> | This Study<br>(CAK 338 /<br>SAD 1483) |
| $\Delta rpoS \Delta cbp$ | $P_{chb}-gfp$, Kan<sup>R</sup>;<br>$\Delta rpoS::Spec^R$; $\Delta cbp::Tm^R$ | A deletion of *lacZ* replaced with $P_{chb}$-GFP transcriptional fusion linked to Kan<sup>R</sup>; a deletion of *cbp* was replaced with Tm<sup>R</sup> and a deletion of *rpoS* replaced with Spec<sup>R</sup> | This Study<br>(CAH 432 /<br>SAD 1486) |
| $\Delta rpoN \Delta cbp$ | $P_{chb}-gfp$, Kan<sup>R</sup>;<br>$\Delta rpoN::Spec^R$; $\Delta cbp::Tm^R$ | A deletion of *lacZ* replaced with $P_{chb}$-GFP transcriptional fusion linked to Kan<sup>R</sup>; a deletion of *cbp* was replaced with Tm<sup>R</sup> and a deletion of *rpoN* replaced with Spec<sup>R</sup> | This Study<br>(CAH 433 /<br>SAD 1487) |
| $\Delta rpoF \Delta cbp$ | $P_{chb}-gfp$, Kan<sup>R</sup>;<br>$\Delta rpoF::Spec^R$; $\Delta cbp::Tm^R$ | A deletion of *lacZ* replaced with $P_{chb}$-GFP transcriptional fusion linked to Kan<sup>R</sup>; a deletion of *cbp* was replaced with Tm<sup>R</sup> and a deletion of *rpoF* replaced with Spec<sup>R</sup> | This Study<br>(CAH 434 /<br>SAD 1488) |
| $\Delta rpoH \Delta cbp$ | $P_{chb}-gfp$, Kan<sup>R</sup>;<br>$\Delta rpoH::Spec^R$; $\Delta cbp::Tm^R$ | A deletion of *lacZ* replaced with $P_{chb}$-GFP transcriptional fusion linked to Kan<sup>R</sup>; a deletion of *cbp* was replaced with Tm<sup>R</sup> and a deletion of *rpoH* replaced with Spec<sup>R</sup> | This Study<br>(CAH 435 /<br>SAD 1489) |
| $\Delta rpoE \Delta cbp$ | $P_{chb}-gfp$, Kan<sup>R</sup>;<br>$\Delta rpoE::Spec^R$; $\Delta cbp::Tm^R$ | A deletion of *lacZ* replaced with $P_{chb}$-GFP transcriptional fusion linked to Kan<sup>R</sup>; a deletion of *cbp* was replaced with Tm<sup>R</sup> and a deletion of *rpoE* replaced with Spec<sup>R</sup> | This Study<br>(CAH 436 /<br>SAD 1490) |
| Ec SBS Ec SlmA | $\Delta lacZ::P_{chb}-GFP$ with Ec<br>SBS, Kan<sup>R</sup>; Ec SlmA;<br>$\Delta VC1807::Spec^R$ | CAK 168 parent with the native copy of SlmA replaced with Ec SlmA; a deletion of VC1807 was replaced with Spec<sup>R</sup> | This Study<br>(CAK 393 /<br>SAD1608) |
| Ec SBS Ec SImA<br>$\Delta cbp$ | $\Delta lacZ::P_{chb}-GFP$ with Ec SBS, Kan<sup>R</sup>; Ec SImA; $\Delta CBP::Spec^R$; | CAK 168 parent with the native copy of SImA replaced with Ec SImA; a deletion of *cbp* was replaced with Spec<sup>R</sup> | This Study (CAK 394 / SAD1609) |
| P<sub>tac</sub>-SImA | $\Delta lacZ::P_{chb}-GFP$, Kan<sup>R</sup>; $\Delta VC1807::P_{tac}$-SImA-3x FLAG, Spec<sup>R</sup> | CAK 007 parent with a deletion of VC1807 replaced with a SImA-3x FLAG fusion linked to the IPTG-inducible P<sub>tac</sub> promoter; P<sub>tac</sub> construct is linked to Spec<sup>R</sup> | This Study (CAK 371 / SAD1614) |
| P<sub>tac</sub>-SImA $\Delta CBP$ | $\Delta lacZ::P_{chb}-GFP$, Kan<sup>R</sup>; $\Delta VC1807::P_{tac}$-SImA-3x FLAG, Spec<sup>R</sup>; $\Delta CBP::Tm^R$ | CAK 007 parent with a deletion of VC1807 replaced with a SImA-3x FLAG fusion linked to the IPTG-inducible P<sub>tac</sub> promoter; P<sub>tac</sub> construct is linked to Spec<sup>R</sup>; a deletion of *cbp* was replaced with Tm<sup>R</sup> | This Study (CAK 372 / SAD1615) |
| P<sub>tac</sub>-SImA F63A | $\Delta lacZ::P_{chb}-GFP$, Kan<sup>R</sup>; $\Delta VC1807::P_{tac}$-SImA F63A-3x FLAG, Spec<sup>R</sup> | Native copy of SImA WT replaced with SImA F63A; deletion of VC1807 replaced with a SImA-3x FLAG fusion linked to the IPTG-inducible P<sub>tac</sub> promoter; P<sub>tac</sub> construct is linked to Spec<sup>R</sup> | This Study (CAK 417 / SAD1616) |
| P<sub>tac</sub>-SImA F63A $\Delta CBP$ | $\Delta lacZ::P_{chb}-GFP$, Kan<sup>R</sup>; $\Delta VC1807::P_{tac}$-SImA F63A-3x FLAG, Spec<sup>R</sup>; $\Delta CBP::Carb^R$ | Native copy of SImA WT replaced with SImA F63A; deletion of VC1807 replaced with a SImA-3x FLAG fusion linked to the IPTG-inducible P<sub>tac</sub> promoter; P<sub>tac</sub> construct is linked to Spec<sup>R</sup>; a deletion of *cbp* was replaced with Carb<sup>R</sup> | This Study (CAK 418 / SAD1617) |
| P<sub>tac</sub>-ChiS | $\Delta lacZ::P_{chb}-GFP$, Kan<sup>R</sup>; $\Delta VCA0692::Ptac$-ChiS, Tm<sup>R</sup> | CAK 007 parent with a deletion of VCA0692 replaced with ChiS linked to the IPTG-inducible P<sub>tac</sub> promoter; P<sub>tac</sub> construct is linked to Tm<sup>R</sup> | This Study (CAK 409 / SAD1619) |
| P<sub>tac</sub>-ChiS $\Delta CBP$ | $\Delta lacZ::P_{chb}-GFP$, Kan<sup>R</sup>; $\Delta VCA0692::Ptac$-ChiS, Tm<sup>R</sup>; $\Delta CBP::Carb^R$ | CAK 007 parent with a deletion of VCA0692 replaced with ChiS linked to the IPTG-inducible P<sub>tac</sub> promoter; P<sub>tac</sub> construct is linked to Tm<sup>R</sup>; a deletion of *cbp* was replaced with Carb<sup>R</sup> | This Study (CAK 410 / SAD1620) |
| P<sub>tac</sub>-SImA P<sub>tac</sub>-ChiS | $\Delta lacZ::P_{chb}-GFP$, Kan<sup>R</sup>; $\Delta VC1807::P_{tac}$-SImA-3x FLAG, Spec<sup>R</sup>; $\Delta VCA0692::Ptac$-ChiS, Tm<sup>R</sup> | CAK 371 parent with a deletion of VCA0692 replaced with ChiS linked to the IPTG-inducible P<sub>tac</sub> promoter; P<sub>tac</sub>-ChiS construct is linked to Tm<sup>R</sup> | This Study (CAK 415 / SAD1621) |
| P<sub>tac</sub>-SImA P<sub>tac</sub>-ChiS $\Delta CBP$ | $\Delta lacZ::P_{chb}-GFP$, Kan<sup>R</sup>; $\Delta VC1807::P_{tac}$-SImA-3x FLAG, Spec<sup>R</sup>; $\Delta VCA0692::Ptac$-ChiS, Tm<sup>R</sup>; $\Delta CBP::Carb^R$ | CAK 371 parent with a deletion of VCA0692 replaced with ChiS linked to the IPTG-inducible P<sub>tac</sub> promoter; P<sub>tac</sub>-ChiS construct is linked to Tm<sup>R</sup>; a deletion of *cbp* was replaced with Carb<sup>R</sup> | This Study (CAK 416 / SAD1622) |
| P<sub>tac</sub>-SImA F63A P<sub>tac</sub>-ChiS | $\Delta lacZ::P_{chb}-GFP$, Kan<sup>R</sup>; $\Delta VC1807::P_{tac}$-SImA F63A-3x FLAG, Spec<sup>R</sup>; | CAK 417 parent with a deletion of VCA0692 replaced with ChiS linked to the IPTG-inducible P<sub>tac</sub> | This Study (CAK 419 / SAD1623) |
| | ΔVCA0692::Ptac-ChiS, Tm<sup>R</sup> | promoter; P<sub>tac</sub>-ChiS construct is linked to Tm<sup>R</sup> | |
| P<sub>tac</sub>-SlmA F63A<br>P<sub>tac</sub>-ChiS ΔCBP | ΔlacZ::P<sub>chb</sub>-GFP, Kan<sup>R</sup>,<br>ΔVC1807::P<sub>tac</sub>-SlmA<br>F63A-3x FLAG, Spec<sup>R</sup>,<br>ΔVCA0692::Ptac-ChiS,<br>Tm<sup>R</sup>; ΔCBP::Carb<sup>R</sup> | CAK 417 parent with a deletion of VCA0692 replaced with ChiS linked to the IPTG-inducible P<sub>tac</sub> promoter; P<sub>tac</sub>-ChiS construct is linked to Tm<sup>R</sup>; a deletion of *cbp* was replaced with Carb<sup>R</sup> | This Study<br>(CAK 420 /<br>SAD1624) |
| **Strains Used for Ec SBS-GFP Repression Reporter Assays** |  |  |  |
| WT SlmA | ΔVC1807::Ec SBS-GFP,<br>Spec<sup>R</sup> | A deletion of VC1807 replaced with consensus sequence of *E. coli* SlmA Binding Sites flanked by -10 and -35 RNA polymerase binding sites fused to GFP and linked to Spec<sup>R</sup> | This Study<br>(CAK 135 /<br>SAD 1427) |
| ΔSlmA | ΔVC1807::Ec SBS-GFP,<br>Spec<sup>R</sup>; ΔslmA::Tm<sup>R</sup> | CAK 135 parent with a deletion of *slmA* replaced with Tm<sup>R</sup> | This Study<br>(CAK 136 /<br>SAD 1428) |
| SlmA T31A | ΔVC1807::Ec SBS-GFP,<br>Spec<sup>R</sup>; SlmA T31A; ΔlacZ<br>LPQEN::Kan<sup>R</sup> | CAK 136 parent with native *slmA* replaced with *slmA* T31A; *lacZ* linked to Kan<sup>R</sup> | This Study<br>(CAK 195 /<br>SAD 1454) |
| SlmA E43A | ΔVC1807::Ec SBS-GFP,<br>Spec<sup>R</sup>; SlmA E43A; ΔlacZ<br>LPQEN::Kan<sup>R</sup> | CAK 136 parent with native *slmA* replaced with *slmA* E43A; *lacZ* linked to Kan<sup>R</sup> | This Study<br>(CAK 244 /<br>SAD 1478) |
| SlmA E43K | ΔVC1807::Ec SBS-GFP,<br>Spec<sup>R</sup>; SlmA E43K; ΔlacZ<br>LPQEN::Kan<sup>R</sup> | CAK 136 parent with native *slmA* replaced with *slmA* E43K; *lacZ* linked to Kan<sup>R</sup> | This Study<br>(CAK 212 /<br>SAD 1461) |
| SlmA F63A | ΔVC1807::Ec SBS-GFP,<br>Spec<sup>R</sup>; SlmA F63A; ΔlacZ<br>LPQEN::Kan<sup>R</sup> | CAK 136 parent with native *slmA* replaced with *slmA* F63A; *lacZ* linked to Kan<sup>R</sup> | This Study<br>(CAK 187 /<br>SAD 1449) |
| SlmA R71D | ΔVC1807::Ec SBS-GFP,<br>Spec<sup>R</sup>; SlmA R71D; ΔlacZ<br>LPQEN::Kan<sup>R</sup> | CAK 136 parent with native *slmA* replaced with *slmA* R71D; *lacZ* linked to Kan<sup>R</sup> | This Study<br>(CAK 188 /<br>SAD 1450) |
| SlmA R173E | ΔVC1807::Ec SBS-GFP,<br>Spec<sup>R</sup>; SlmA R173E;<br>ΔlacZ LPQEN::Kan<sup>R</sup> | CAK 136 parent with native *slmA* replaced with *slmA* R173E; *lacZ* linked to Kan<sup>R</sup> | This Study<br>(CAK 189 /<br>SAD 1451) |
| **Strains Used for Microscopy** |  |  |  |
| WT SlmA | ΔlacZ::P<sub>tac</sub>-SlmA, Spec<sup>R</sup> | *lacZ* replaced with SlmA WT under the control of an IPTG inducible P<sub>tac</sub> promoter (derived from a Spec<sup>R</sup> Tn10 fragment containing LacI, Spec<sup>R</sup>, and an outward reading P<sub>tac</sub> promoter) | This Study<br>(CAK 137 /<br>SAD 1429) |
| SlmA T31A | ΔlacZ::P<sub>tac</sub>-SlmA, Spec<sup>R</sup> | *lacZ* replaced with SlmA T31A under the control of an IPTG inducible P<sub>tac</sub> promoter (derived from a Spec<sup>R</sup> Tn10 fragment containing LacI, Spec<sup>R</sup>, and an outward reading P<sub>tac</sub> promoter) | This Study<br>(CAK 151 /<br>SAD 1432) |
| SlmA E43A | ΔlacZ::P<sub>tac</sub>-SlmA, Spec<sup>R</sup> | *lacZ* replaced with SlmA E43A under the control of an IPTG inducible P<sub>tac</sub> promoter (derived from a Spec<sup>R</sup> Tn10 fragment containing LacI, Spec<sup>R</sup>, and an outward reading P<sub>tac</sub> promoter) | This Study<br>(CAK 152 /<br>SAD 1433) |
| SlmA E43K | $\Delta lacZ::P_{tac}$-SlmA, Spec<sup>R</sup> | *lacZ* replaced with SlmA E43K under the control of an IPTG inducible $P_{tac}$ promoter (derived from a Spec<sup>R</sup> Tn10 fragment containing LacI, Spec<sup>R</sup>, and an outward reading $P_{tac}$ promoter) | This Study (CAK 341 / SAD 1484) |
| SlmA F63A | $\Delta lacZ::P_{tac}$-SlmA, Spec<sup>R</sup> | *lacZ* replaced with SlmA F63A under the control of an IPTG inducible $P_{tac}$ promoter (derived from a Spec<sup>R</sup> Tn10 fragment containing LacI, Spec<sup>R</sup>, and an outward reading $P_{tac}$ promoter) | This Study (CAK 153 / SAD 1434) |
| SlmA R71D | $\Delta lacZ::P_{tac}$-SlmA, Spec<sup>R</sup> | *lacZ* replaced with SlmA R71D under the control of an IPTG inducible $P_{tac}$ promoter (derived from a Spec<sup>R</sup> Tn10 fragment containing LacI, Spec<sup>R</sup>, and an outward reading $P_{tac}$ promoter) | This Study (CAK 154 / SAD 1435) |
| SlmA R173E | $\Delta lacZ::P_{tac}$-SlmA, Spec<sup>R</sup> | *lacZ* replaced with SlmA R173E under the control of an IPTG inducible $P_{tac}$ promoter (derived from a Spec<sup>R</sup> Tn10 fragment containing LacI, Spec<sup>R</sup>, and an outward reading $P_{tac}$ promoter) | This Study (CAK 342 / SAD 1485) |
| **Strains Used for SlmA Overexpression and Purification** |  |  |  |
|  | E. coli BL21-DE3 harboring pHisTev w / WT SlmA | WT SlmA overexpression strain – SlmA cloned into NdeI / BamHI sites | This Study (CAK 110 / SAD 1414) |
|  | E. coli BL21-DE3 harboring pHisTev + SlmA T31A | SlmA T31A overexpression strain – vector generated using single primer site directed mutagenesis | This Study (CAK 161 / SAD 1436) |
|  | E. coli BL21-DE3 harboring pHisTev + SlmA E43A | SlmA E43A overexpression strain – vector generated using single primer site directed mutagenesis | This Study (CAK 162 / SAD 1437) |
|  | E. coli BL21-DE3 harboring pHisTev + SlmA E43K | SlmA E43K overexpression strain – vector generated using single primer site directed mutagenesis | This study (CAK 268 / SAD 1479) |
| **Strains Used for FLAG Western Blots** |  |  |  |
| SlmA WT-3x FLAG | $\Delta lacZ::P_{chb}$-GFP, Kan<sup>R</sup>; SlmA WT-3x FLAG; $\Delta CBP::Spec^R$ | CAK 024 parent with SlmA WT-3x FLAG knocked back in at native locus; deletion of *cbp* replaced with Spec<sup>R</sup> | This study (CAK 426 / SAD 1601) |
| SlmA T31A-3x FLAG | $\Delta lacZ::P_{chb}$-GFP, Kan<sup>R</sup>; SlmA T31A-3x FLAG; $\Delta CBP::Spec^R$ | CAK 024 parent with SlmA T31A-3x FLAG knocked back in at native locus; deletion of *cbp* replaced with Spec<sup>R</sup> | This study (CAK 380 / SAD 1602) |
| SlmA E43A-3x FLAG | $\Delta lacZ::P_{chb}$-GFP, Kan<sup>R</sup>; SlmA E43A-3x FLAG; $\Delta CBP::Spec^R$ | CAK 024 parent with SlmA E43A-3x FLAG knocked back in at native locus; deletion of *cbp* replaced with Spec<sup>R</sup> | This study (CAK 382 / SAD 1603) |
| SlmA E43K-3x FLAG | $\Delta lacZ::P_{chb}$-GFP, Kan<sup>R</sup>; SlmA E43K-3x FLAG; $\Delta CBP::Spec^R$ | CAK 024 parent with SlmA E43K-3x FLAG knocked back in at native locus; deletion of *cbp* replaced with Spec<sup>R</sup> | This study (CAK 384 / SAD 1604) |
| SlmA F63A-3x FLAG | $\Delta lacZ::P_{chb}$-GFP, Kan<sup>R</sup>; SlmA F63A-3x FLAG; $\Delta CBP::Spec^R$ | CAK 024 parent with SlmA F63A-3x FLAG knocked back in at native locus; deletion of *cbp* replaced with Spec<sup>R</sup> | This study (CAK 386 / SAD 1605) |
| SlmA R71D-3x FLAG | $\Delta lacZ::P_{chb}$-GFP, Kan<sup>R</sup>; SlmA R71D-3x FLAG; $\Delta CBP::Spec^R$ | CAK 024 parent with SlmA R71D-3x FLAG knocked back in at native locus; deletion of *cbp* replaced with Spec<sup>R</sup> | This study (CAK 388 / SAD 1606) |
| SlmA R173E-3x FLAG | $\Delta lacZ::P_{chb}$-GFP, Kan<sup>R</sup>; SlmA R173E-3x FLAG; $\Delta CBP::Spec^R$ | CAK 024 parent with SlmA R173E-3x FLAG knocked back in at native locus; deletion of *cbp* replaced with Spec<sup>R</sup> | This study (CAK 390 / SAD 1607) |
| Ptac-ChiS-FLAG | $\Delta lacZ::P_{chb}$-GFP, Kan<sup>R</sup>; $\Delta VCA0692::Ptac$-ChiS-1x FLAG, Tm<sup>R</sup>; $\Delta ChiS::Spec^R$; $\Delta CBP::Carb^R$ | CAK 007 parent with a deletion of VCA0692 replaced with a ChiS-1x FLAG fusion linked to the IPTG-inducible P<sub>tac</sub> promoter; P<sub>tac</sub>-ChiS construct is linked to Tm<sup>R</sup>; deletion of *chiS* replaced with Spec<sup>R</sup> and a deletion of *cbp* replaced with Carb<sup>R</sup> | This study (CAK 406 / SAD 1618) |
| ChiS-FLAG | $\Delta lacZ::P_{chb}$-GFP, Kan<sup>R</sup>; $\Delta ChiS::ChiS$-1x FLAG; $\Delta CBP::Tm^R$ | CAK 007 parent with native ChiS replaced with a ChiS-1x FLAG fusion; deletion of CBP replaced with Tm<sup>R</sup> | This study (CAK 374 / SAD 1625) |
| **Strains Used for mCherry assays** |  |  |  |
| WT | $\Delta lacZ::P_{chb}$-mCherry, Kan<sup>R</sup> | A deletion of *lacZ* replaced with P<sub>chb</sub>-mCherry transcriptional fusion linked to Kan<sup>R</sup> | This study (CAK 346 / SAD 1610) |
| $\Delta cbp$ | $\Delta lacZ::P_{chb}$-mCherry, Kan<sup>R</sup>; $\Delta CBP::Spec^R$ | CAK 346 parent with a deletion of *cbp* replaced with Spec<sup>R</sup> | This study (CAK 347 / SAD 1611) |
| SlmA-3x FLAG | $\Delta lacZ::P_{chb}$-mCherry, Kan<sup>R</sup>; $\Delta SlmA::SlmA$-3x FLAG; $\Delta VC1807::Spec^R$ | CAK 346 parent with the native copy of SlmA replaced with SlmA-3x FLAG; deletion of VC1807 replaced with Spec<sup>R</sup> | This study (CAK 358 / SAD 1612) |
| SlmA-3x FLAG $\Delta cbp$ | $\Delta lacZ::P_{chb}$-mCherry, Kan<sup>R</sup>; $\Delta SlmA::SlmA$-3x FLAG; $\Delta CBP::Spec^R$ | A deletion of *lacZ* replaced with P<sub>chb</sub>-mCherry transcriptional fusion linked to Kan<sup>R</sup>; native copy of SlmA replaced with SlmA-3x FLAG; deletion of CBP replaced with Spec<sup>R</sup> | This study (CAK 359 / SAD 1613) |

**Table S1.**
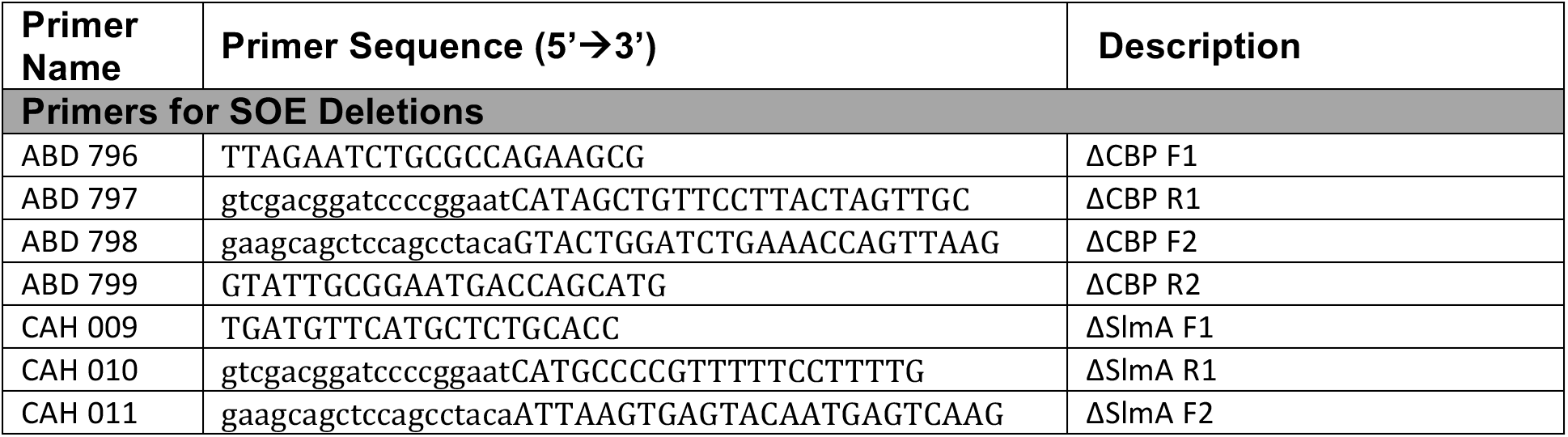
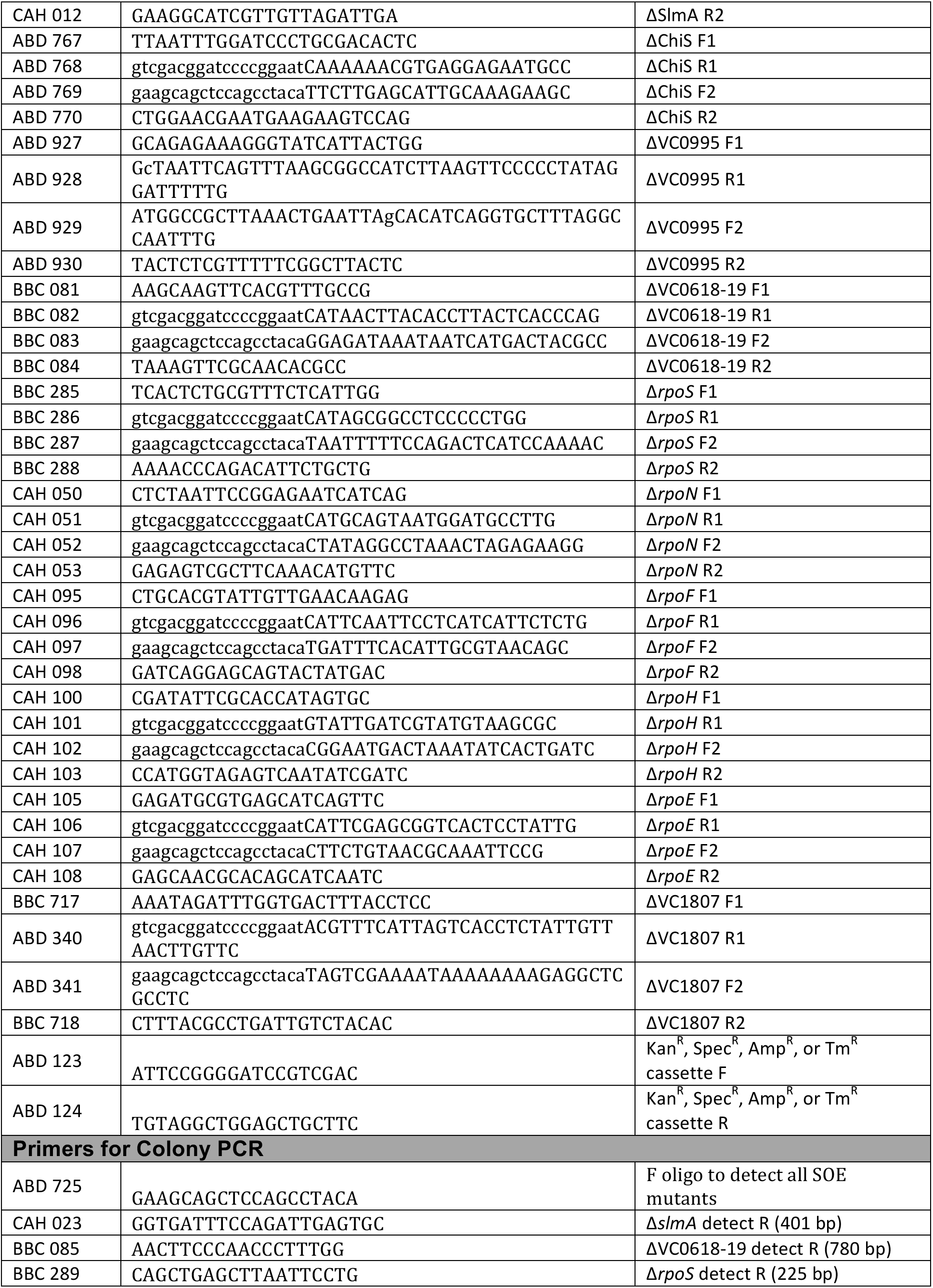
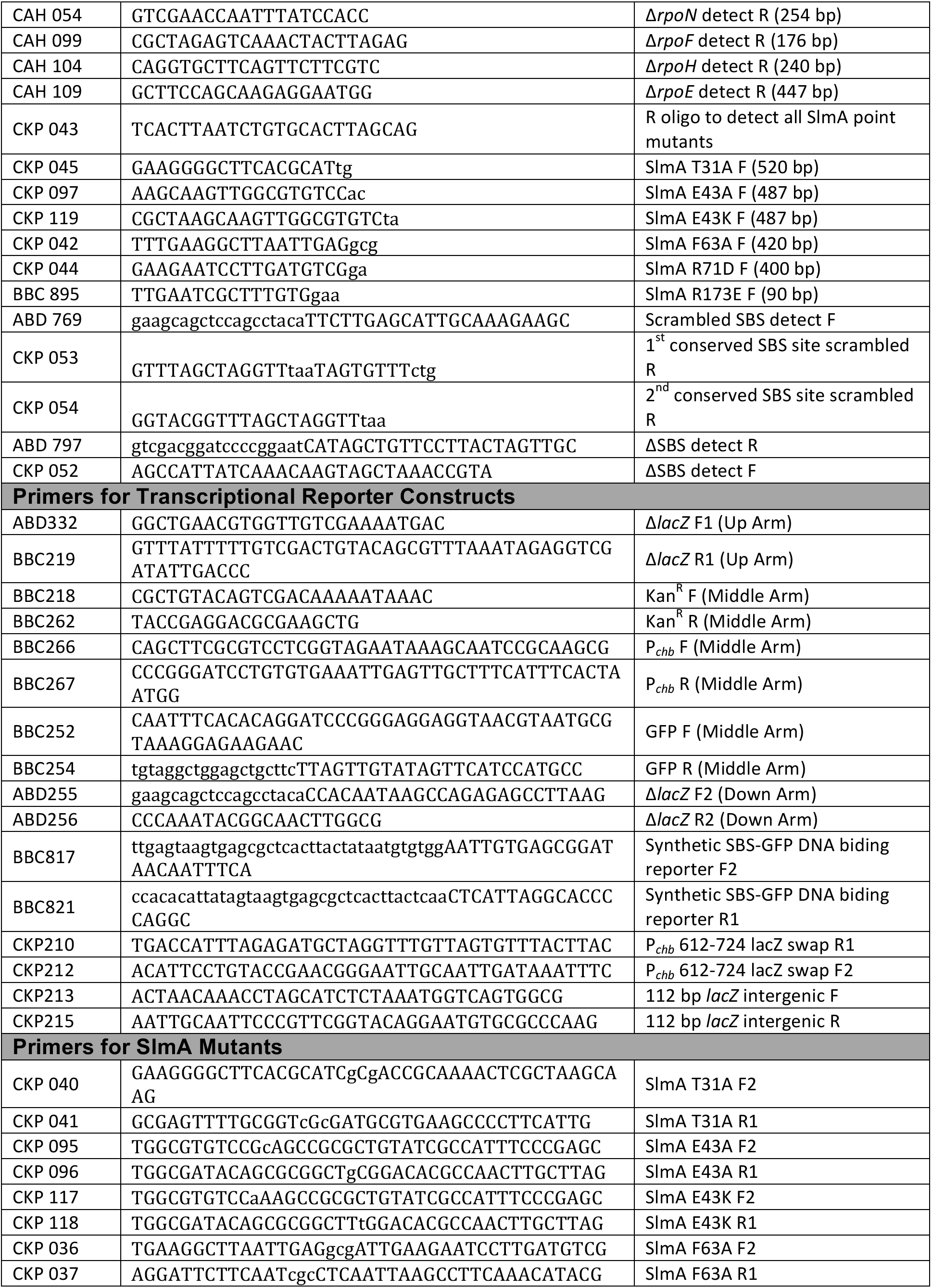
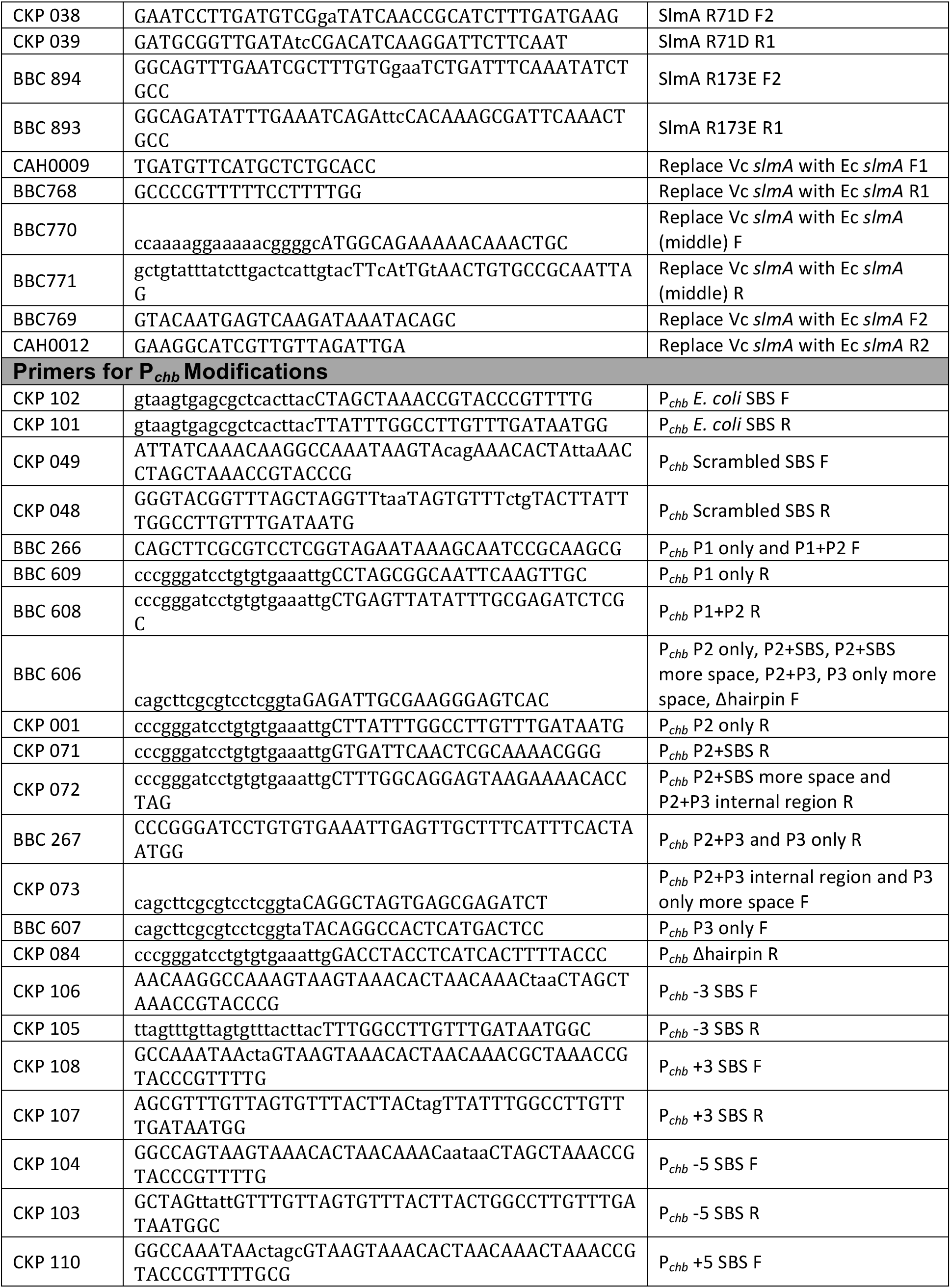
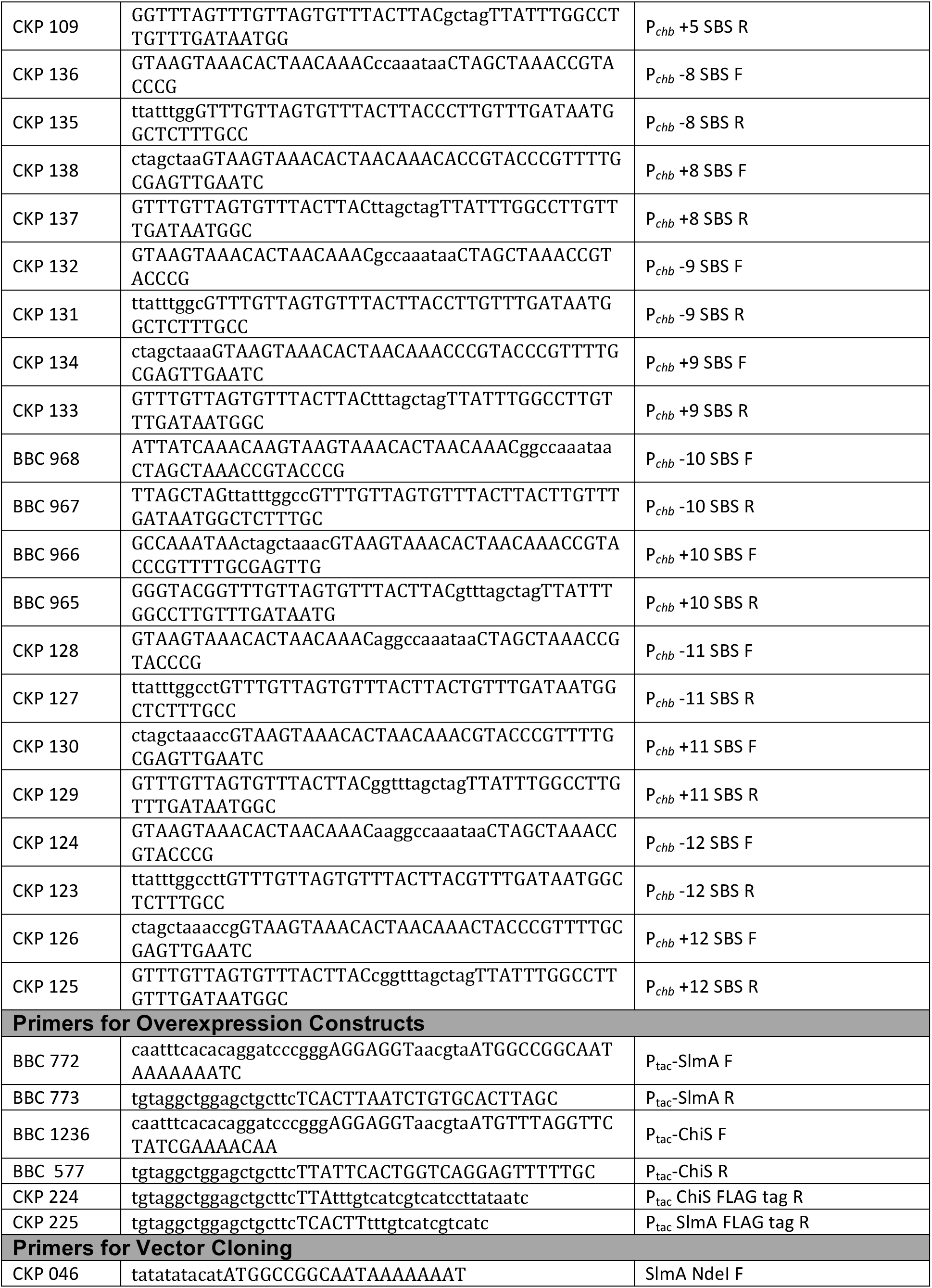
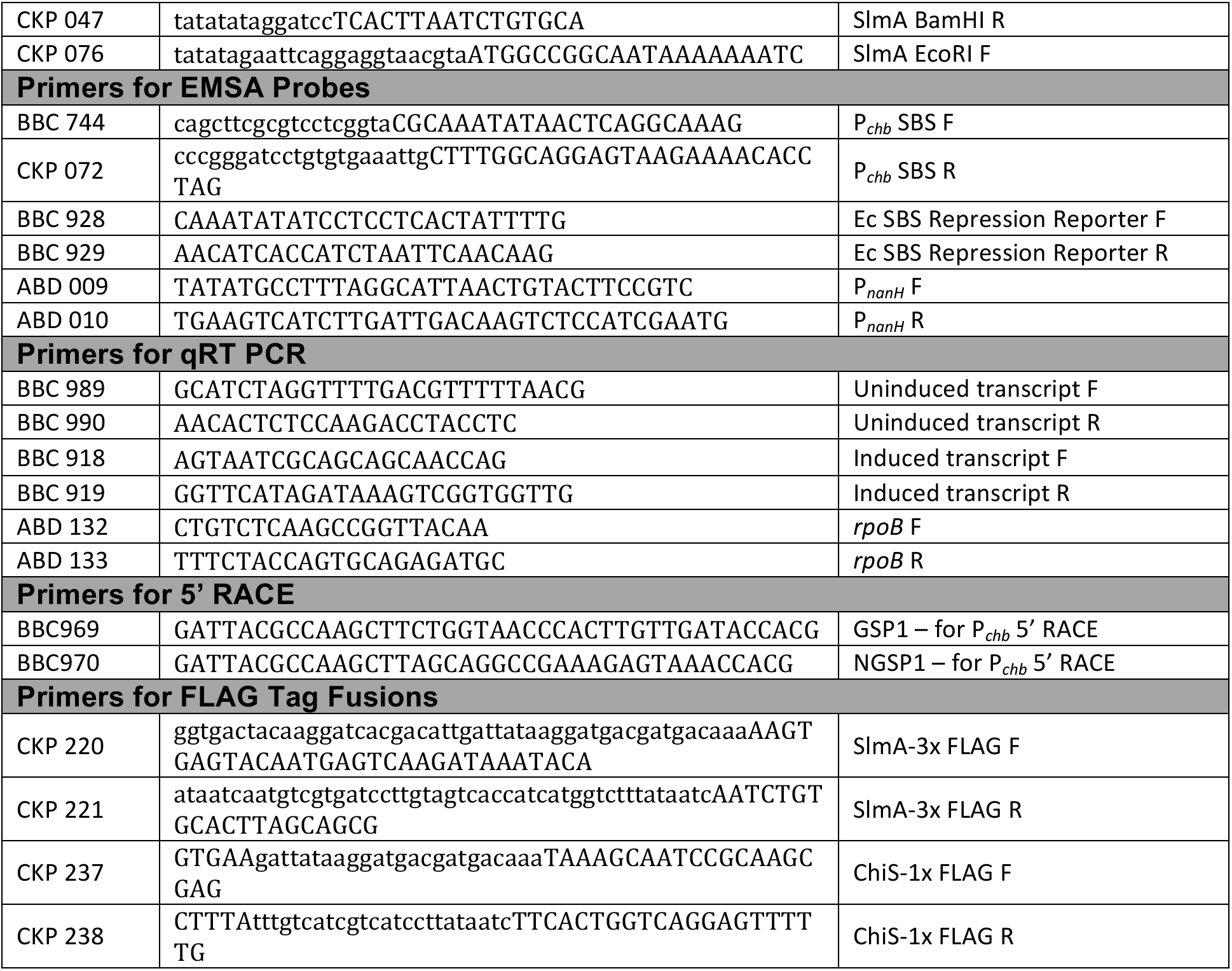
-Primers used in this study

